# Early-life gut microbiota assembly patterns are conserved between laboratory and wild mice

**DOI:** 10.1101/2023.10.18.562850

**Authors:** Eveliina Hanski, Aura Raulo, Sarah C L Knowles

**Author notes:** **Competing interests** Authors declare no competing interests.

## Abstract

Assembly of the mammalian gut microbiota during early life is known to shape key aspects of organismal development, including immunity, metabolism and behaviour. While house mice (*Mus musculus*) are the major laboratory model organism for gut microbiota research, their artificial lab-based lifestyle could fundamentally alter ecological processes of microbiota assembly and dynamics, in ways that affect their usefulness as a model system. To examine this, here we directly compared patterns of gut microbiota assembly in house mice from the lab and from the wild, making use of a tractable, individually-marked wild population where we could examine patterns of gut microbiota assembly during early life. Despite lab and wild mice harbouring taxonomically distinct communities, we identify striking similarities in multiple patterns of their gut microbiota assembly. Specifically, age-related changes in both alpha and beta diversity, as well as the abundance of predominant phyla and aerotolerance of the microbiota followed parallel trajectories in both settings. These results suggest some degree of intrinsic programme in gut microbiota assembly that transcends variation in taxonomic profiles, and the genetic and environmental background of the host. They further support the notion that despite their artificial environment, lab mice can provide meaningful insights into natural microbiota ecological dynamics in early life and their interplay with host development.

## Introduction

The mammalian gut microbiota has diverse impacts on host biology with key roles modulating processes ranging from metabolism to behaviour^1–3^. The composition and diversity of this internal ecosystem varies between individuals, populations, and species and also within individuals over time^4–6^. The gut microbiota matures in early life, during which marked changes in composition and diversity occur^7–9^. This maturation is guided by both ecological interactions within the microbial community^10^ and selective forces imposed by the hosts’ physiology as it undergoes key developmental changes including development of the immune and central nervous systems^11,12^. Perturbation of the gut microbiota during this critical period has been linked to altered functioning of the immune system in early life^13,14^, indicating an active role for the gut microbiota in host development. Understanding the patterns and drivers of this microbiota maturation process is therefore essential for a comprehensive understanding of host development.

At birth, the gut is first colonised by microbes from the mother (through birth and breastfeeding) and the environment, after which the microbiota matures following a stereotypical pattern in which the community becomes more diverse within individuals but more homogenous across individuals^15,16^. In humans, it is well-established that environmental context can alter the microbiota’s maturation trajectory at various stages, with potentially long-lasting consequences^14,17,18^. Delivery mode affects the transmission of maternal microbes at birth, with C-section delivery predisposing infants to greater colonisation by microbes with pathogenic potential from the hospital environment over specialist beneficial symbionts from the mother^7,15,19^. Subsequently, gut microbiota maturation can be altered by care practices such as feeding mode (breastfeeding/formula) and the timing of breastfeeding cessation^7,19,20^, which affect both the transmission landscape and selective forces shaping the developing microbiota. Microbiota maturation is also known to be sensitive to environmental influences later in childhood^21–23^.

The gut microbiota’s stereotypical trajectory of maturation has been well-characterised in humans, but in other mammals and especially in wild contexts, it is far less well-known. Studies of gut microbiota assembly in wild primates have reported somewhat conflicting patterns, with research on gelada baboons finding patterns that largely reflect those in humans^24,25^, while a study in wild chimpanzees reported declining microbiota richness during early life^26^. This suggests the broad patterns of microbiota assembly observed in humans may not be universal among mammals, and may vary with host genetic background or environmental context. However, to our knowledge there are currently no studies of mammals other than humans which directly compare patterns of gut microbiota assembly across populations or environmental contexts, such that we currently do not know the extent to which context shapes gut microbiota assembly patterns for a given host species.

Whether or not gut microbiota maturation patterns are consistent or variable across contexts is a particularly pertinent question for laboratory-reared model organisms. The laboratory mouse is the major model species for fundamental and preclinical research on mammals, and aspects of its gut microbiota maturation have been studied in several laboratory strains^27–34^. In line with patterns detected in other mammals, existing studies suggest that as lab mice mature, the gut microbiota increases in diversity within individuals while becoming more similar across individuals^27,28,33,34^.

Importantly, the dynamics of gut microbiota assembly in wild house mice have not yet been characterised, such that it remains unknown whether gut microbiota assembly in laboratory mice is representative of that occurring in natural populations. In the wild, many factors that could affect both microbial colonisation and subsequent within-host selective processes will differ from and be far more variable than in a laboratory. These include which specific microbes are present, how and when mice are exposed to them, as well as diet and host genetics. Given its model species status and the known influence of the developing gut microbiota on so many aspects of physiology, understanding how well lab mouse gut microbiota assembly dynamics reflect those occurring naturally in wild mouse populations, is an important goal.

Here, using standardised microbiota profiling methods we directly compare the dynamics of gut microbiota assembly in young laboratory mice of the most commonly used strain (C57BL/6) and completely wild house mice sampled from the island of Skokholm, UK. In contrast to inbred lab mice living under consistently stable environmental conditions, this wild population are outbred and exposed to highly variable environmental conditions. Given the gut microbiota is thought to play a key role in developmental processes that should be conserved across these contexts, we hypothesised that despite some differentiation in community composition between lab and wild mice, some broad trends of microbiota assembly would be conserved.

## Materials and Methods

### Sample collection

Faecal samples were collected from 39 individual C57BL/6 laboratory mice (*Mus musculus*) aged between 7 and 124 days (mean 39), in October 2021 at the Biomedical Services Building, Oxford, UK. The colony originated from Jackson Laboratories but has been bred at the facility for the last 17 years. Mice were housed under specific-pathogen-free (SPF) conditions, where the following pathogenic taxa were excluded: *Helicobacter* species, *Pasteurella pneumotropica,* beta-haemolytic streptococci*, Streptococcus pneumoniae, Citrobacter rodentium, Corynebacterium kutscheri, Salmonella* species, and *Streptobacillus moniliformis*. Mice were not subject to any interventions before or during sample collection. Mice were separated from dams at 19–22 days of age. Faecal samples were collected by placing mice on a sterile surface. Pellets were collected with sterile forceps, preserved in DNA/RNA Shield (Zymo Research, USA) and stored at -80°C until DNA extraction (≤12 months).

Wild house mice (*Mus musculus domesticus*) were live-trapped on Skokholm Island (UK; Figure S1) in September–October 2019 and August–September 2020. House mice have been living relatively independent from humans on Skokholm since their introduction around the turn of the 20th century, and there are no other small mammals on the island^35,36^. Trapping was carried out in two sampling sites on the island (Fig. S1) using small Sherman traps provisioned with peanuts, non-absorbent cotton wool for bedding, and with a spray of sesame oil outside the trap used as a lure. On each trapping night, 150 traps were set at dusk and checked at dawn at one of the sampling sites. Visited traps (where signs of a mouse were detected, whether captured or not) were washed and sterilised with 20% bleach solution before re-use. Captured mice were tagged with a subcutaneous passive integrated transponder (PIT) tag for permanent identification, or identified through PIT tag detection upon recapture. All captures were therefore individually identified, sexed, scored for reproductive status, and weighed to the nearest 0.1g before release at their trapping point. Sex was determined based on measurement of anogenital distance. Faecal pellets were collected from traps, and preserved in DNA/RNA Shield. Samples were stored at -20°C during fieldwork (≤6 weeks), after which they were transported frozen to the laboratory and stored at -80°C until DNA extraction (≤17 months).

Since wild-caught mice cannot be precisely aged, we used body mass as a proxy for age, as among young individuals has been shown to predict age relatively accurately in both wild and laboratory mice (Fig. S2)^37–40^. Female mice showing signs of pregnancy (bulging central body; *n*=15) were excluded, as these were associated with heavier body mass. We also excluded female mice ≥25g (*n*=65), with the aim of excluding pregnant females that may not have been recognised as such. Applying these criteria resulted in a set of 474 faecal samples from 235 individual wild mice (1–10 samples each).

### DNA extraction, library preparation, and sequencing

DNA extraction was performed using the ZymoBIOMICS DNA MiniPrep Kit (spin-column format) according to manufacturer’s instructions (Zymo Research, USA). Samples were randomised into 53 extraction batches of up to 23 samples. A negative extraction control (DNAse-free H_2_O) was included in each extraction batch at a varying position with the exception of one extraction batch, in which a negative control was not included. The microbiota of each faecal sample was characterised using amplicon sequencing, targeting the V4–V5 region of the 16S rRNA gene using primers 515F and 926R^41,42^. Library preparation was conducted in 16 plates of up to 95 samples. Amplicons were sequenced in four sequencing runs using the Illumina MiSeq platform (Reagent kit v3, 2x300 bp chemistry). Library preparation and sequencing was conducted at the Integrated Microbiome Resource, using the protocol described in Comeau et al. (2017)^43^. Each sequencing run included a PCR reaction negative control and a separate negative control for sequencing. All extraction controls were sequenced intermixed with true samples.

### Data processing

Data processing and subsequent analyses were carried out using R v4.1.2. Demultiplexed sequencing reads were processed through the DADA2 pipeline^44^ to infer amplicon sequence variants (ASVs) and assign taxonomy using the SILVA rRNA database v138.1. A phylogenetic tree was built using packages *DECIPHER* and *phangorn*. The package *iNEXT*^45^ was used to generate sample completeness and rarefaction curves separately for lab and wild mouse datasets. Based on these curves, lab and wild mouse samples with a read depth below 7,500 and 4,000, respectively, (where the curves plateaued) were excluded from further analysis. *iNEXT* was further used to measure asymptotic ASV richness and Shannon diversity. In each of the lab and wild mouse datasets separately, singleton and doubleton ASVs, as well as those assigned as mitochondria or chloroplast sequences were removed. ASV counts were normalised to proportional abundance using package *phyloseq*^46^. Information on bacterial aerotolerance was extracted from Bergey’s Manual of Systematics of Archaea and Bacteria or additional references when this information was not available in the manual (Table S1). Aerotolerance was determined based on genus-level taxonomy, unless aerotolerance information was not available or genus-level taxonomy not assigned, in which case family-level information was inspected and used if all genera within a family were stated to have the same aerotolerance. Bacteria were categorised as *aerotolerant* or *obligate anaerobe*. If a given genus included both obligate anaerobes and aerotolerants, a Nucleotide BLAST search was conducted for the associated ASVs and if a 100% species identity match was found, aerotolerance category was determined based on this species.

A total of 41 unique ASVs were detected across all negative DNA extraction and library preparation controls (*n*=66), with a maximum read count of 363 for any given control, with the exception of one library preparation control. This control contained 255 unique ASVs and 29,300 reads (mean read count of biological samples was 29,235). All extraction controls (*n*=5) from this control’s 96-well plate had <20 reads each, indicating the entire plate was not contaminated during library preparation (negative controls were included on the plate in a randomised order, thus several rows and columns had negative controls). Instead, the most likely explanation is that a biological sample was mistakenly pipetted into the control well in addition to its designated well. As all extraction controls on this plate showed very low read counts indicating no systemic contamination, we retained this plate of samples in our analyses.

We tested for the presence of any potential contaminants using the R package *decontam*^47^ using the ‘prevalence’ method, in which the prevalence of each sequence in biological samples is compared to that in negative controls (excluding the library preparation control that was contaminated with a biological sample, see above). A sequence was considered a contaminant if it reached a probability of 0.1 in the Fisher’s exact test used in *decontam*. 31 contaminants were identified and filtered from the data before subsequent analysis.

### Functional profiles

Functional pathways were predicted from the 16S rRNA data using PICRUSt2, pipeline v 2.5.0^48^. Read counts of pathways were normalised to proportional abundance. Functional profiles were characterised at pathway-level only, as functional profiles characterised at category level varied depending which database used to assign functional categories (MetaCyc vs KEGG ortholog). Data presented is based on MetaCyc pathways.

### Analysis

Principal coordinate analysis was used to investigate source and age effects on microbiota composition. To explore different aspects of compositional microbiota variation in these analyses we used four different measures of beta diversity: Jaccard, Aitchison, unweighted UniFrac and weighted UniFrac distances. Beta diversity metrics were calculated with packages *vegan*^49^ and *microbiome*^50^. Statistical differences in alpha and beta diversity were tested with Wilcoxon rank sum tests (with 1,000 permutations for beta diversity analyses).

Predictors of beta diversity variation were investigated using marginal permutational multivariate analysis of variance (PERMANOVA) run using the adonis2 function in *vegan*. Quadratic plateau models run in package *easynls*^51^ were used to assess how microbiota alpha diversity, and gut microbiota maturity (distance to a juvenile reference gut microbiota) changed over time. To enable direct comparison of age-related patterns across systems in these models, changes in microbiota traits over time in these models and their associated figures are presented as a function of body mass. Although individual body mass measurements were not available for the laboratory mouse dataset, age is tightly correlated with mass among juvenile lab and wild house mice^37,34^, such that we used estimated body mass values for lab mice from published age-mass relationships (Fig. S2). For mice ≤21 days of age, body mass was estimated using mean body mass for each unit of age reported in Figure 3 in Spangenberg et al. (2014)^37^ (no separation based on sex). For mice >21 days of age, body mass was estimated separately for male and female mice by using the mean body mass for each unit of age reported in ‘*Body weight information for C57BL/6J*’ table on The Jackson Laboratory website^38^.

The reference community of early life microbiota was estimated separately for laboratory and wild mice. This estimation was based on the average abundance observed in mice that were under 15 days of age (estimated age for wild mice). This corresponds to an estimated body mass ≤6.5g for lab mice, and a measured body mass ≤7.9g for wild mice^37,40^. The choice of age cut-off (14 days) was made by identifying the youngest age at which there were at least six samples available in both laboratory and wild mouse data. With this cut-off, 6 lab mice (body mass range 3.5–6.5g, mean 5.4g) and 16 wild mice (body mass range 5.4–7.8g, mean 6.9g) were included in estimating the reference communities. Relative abundances of taxonomic groups or phenotypic subsets of bacteria were modelled and plotted as a function of body mass using locally weighted scatterplot smoothing (LOESS) regression in package *ggplot2*^52^.

## Results

A total of 8,384 ASVs were recovered across lab and wild mice. Taxonomic assignment rates were comparable across the two systems, indicating lab and wild mice have similar numbers of identifiable bacterial genera (89.9% and 84.3% ASVs assigned to family, 50.0% and 51.9% assigned to genus, for lab and wild mice respectively).

### Laboratory and wild mice have distinct gut microbiotas

To facilitate statistical comparisons between the two settings, we compared gut microbiota of lab and wild mice using a randomly selected cross-sectional subset of the data (39 samples from each system). In this subset, only 12.6% ASVs (369 of 2,920) were detected in both lab and wild mice. However, these shared taxa were common and abundant in both lab and wild mice, comprising on average 41.2% of the wild mouse microbiota and 43.1% of the lab mouse microbiota (standard deviation 9.1% and 20.4%, respectively). Wild mice had higher alpha diversity, both in terms of estimated ASV richness and Shannon diversity (Wilcoxon rank sum tests, *p*<0.001 for both, Fig. 1A–B). Lab and wild mouse gut microbiotas were broadly similar at the phylum level, but showed clear differences in composition at lower taxonomic levels (Fig. 1C–D). Compositionally, samples clustered by source using both non-phylogenetic and phylogenetic distance metrics and using various levels of taxonomic resolution (Fig. 1E, Fig. S3). In line, source explained a considerable amount of compositional variation in the dataset overall (11.8% in Jaccard dissimilarity, PERMANOVA, *p*=0.001; beta dispersion, *F*=0.3758, *p*=0.563; 18.2% in Aitchison dissimilarity, *p*=0.001; beta dispersion, *F*=68.172, *p*=0.001). Wild mouse individuals were much more variable in microbiota composition than lab mice, although lab–wild sample pairs had the highest dissimilarity (Fig. 1D, all permutational Wilcoxon rank sum tests *p*<0.001).

**Figure 1.**
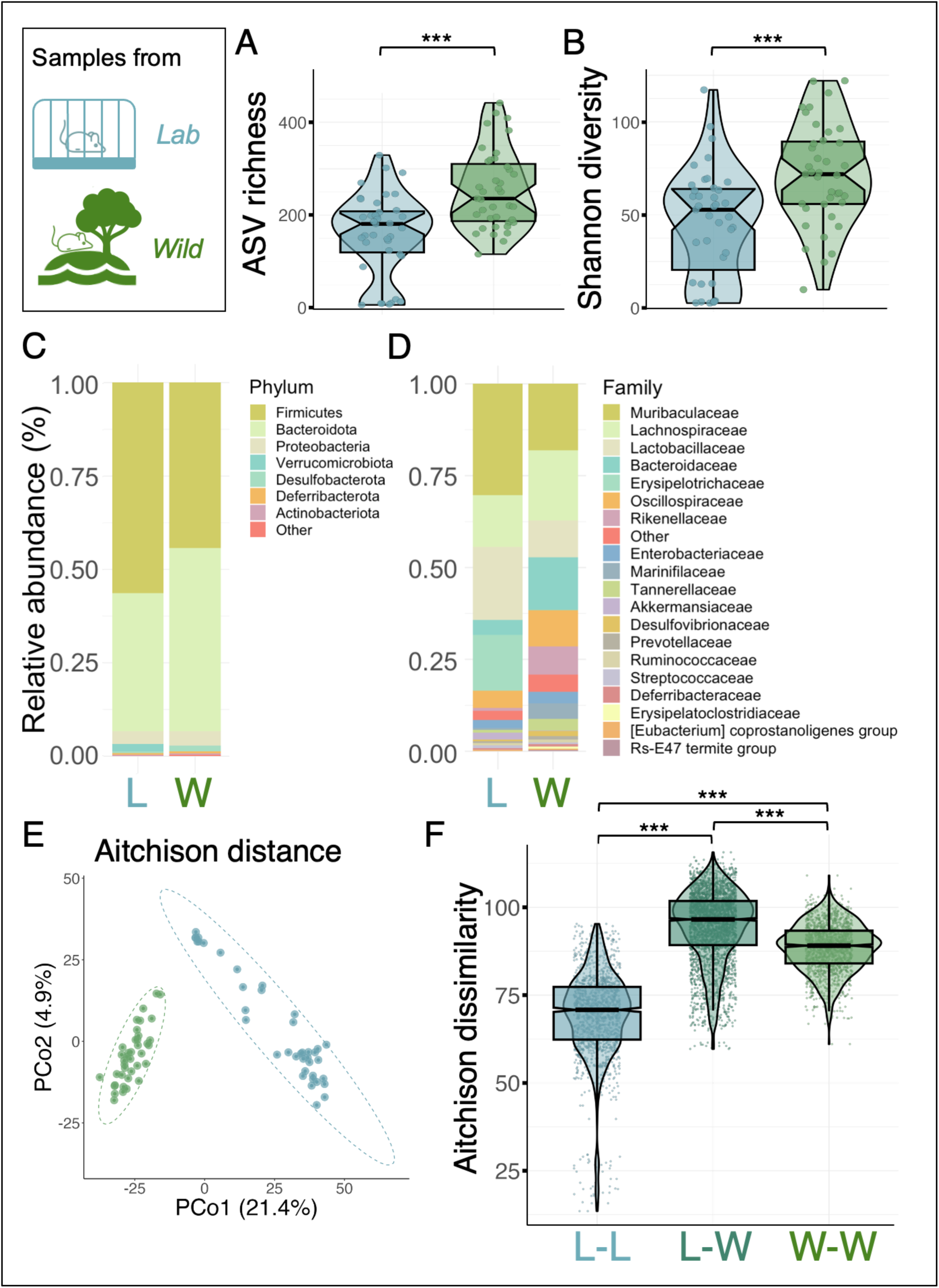
Comparison of laboratory and wild mouse gut microbiotas using faecal samples from 39 laboratory and 39 wild mice (1 randomly selected sample per mouse). (**A–B**) Alpha diversity; (**A**) asymptotic ASV richness and (**B**) asymptotic Shannon diversity. Statistical differences between lab and wild mice were tested with Wilcoxon rank sum tests (***; *p*<0.001). (**C–D**) Mean relative abundance of bacterial (**C**) phyla and (**D**) families in laboratory (L) and wild (W) mice. (**E**) Principal coordinates analysis (PCoA) on Aitchison distance. (**F**) Pairwise comparison on gut microbial dissimilarity on Aitchison distance in lab–lab (L–L), lab–wild (L–W) and wild–wild (W–W) sample pairs. Statistical differences between lab and wild mice were tested with permutational Wilcoxon rank sum tests (***; *p*<0.001). Colour in figures A–B and E–F indicates sample source: *blue* = laboratory, *green* = wild, *teal* = lab/wild.

### Conserved taxon-independent gut microbiota assembly patterns

In the absence of known age of the wild mice included in the study, we used body mass as a proxy of age. Previous studies have shown that in the first 30 days of life the relationship between age and body mass is very similar in lab and wild *Mus musculus*^37–40^ (Fig. S2). Body mass was strongly associated with alpha diversity in both lab and wild mice, with ASV richness increasing to a plateau in both systems. Parameters from quadratic plateau models indicated richness plateaud at 281 ASVs in adulthood among wild mice, and at 213 ASVs in lab mice (maximum values of y). In lab mice, the plateau in ASV richness was estimated to occur at a body mass of 13 grams (Fig. 2A), or 27 days of age (Fig. S2). The plateau in ASV richness was more subtle in wild mice, and occurred a bit later around 21 grams (Fig. 2B).

**Figure 2.**
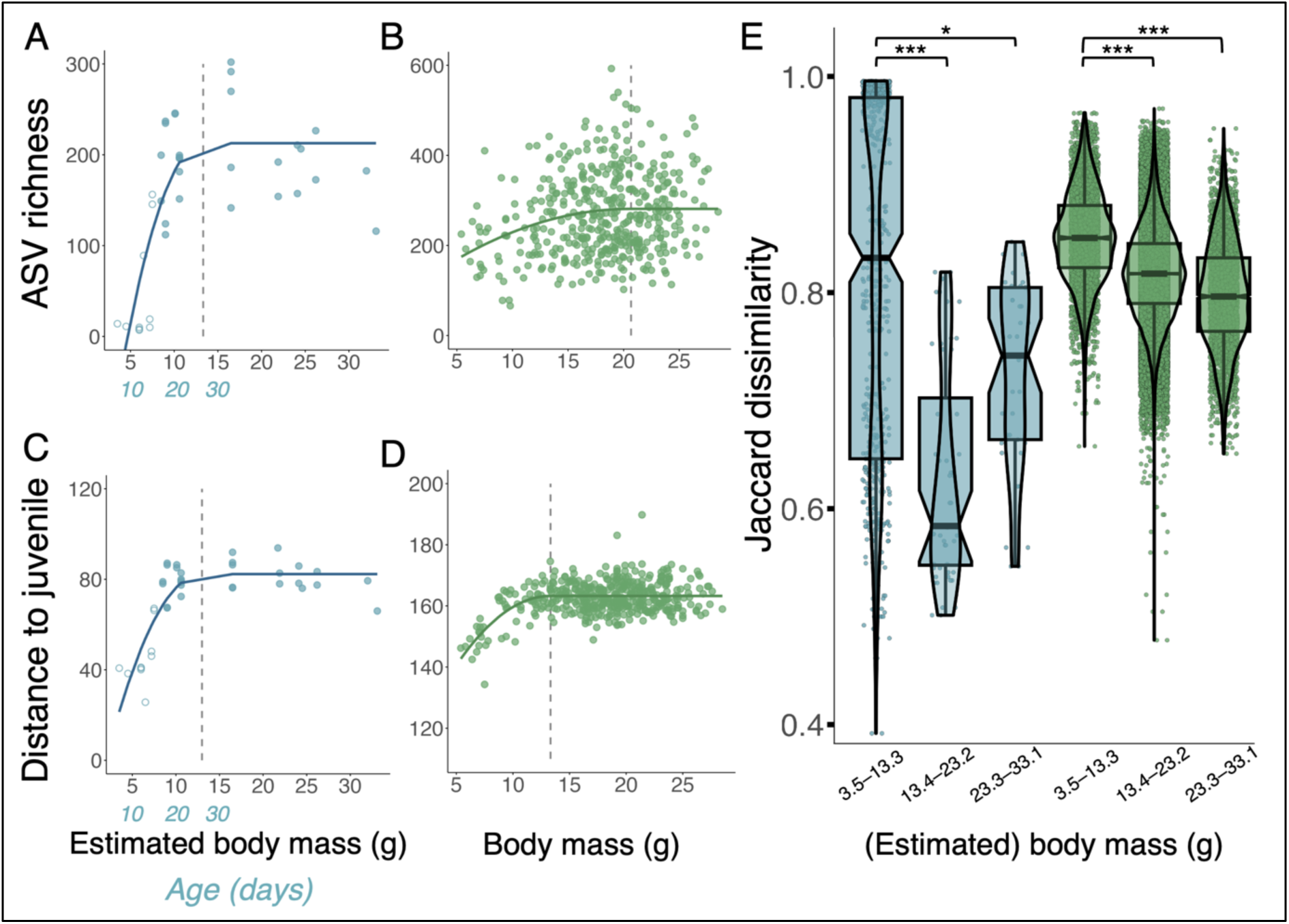
Taxon-independent trends in laboratory (*blue*) and wild house mouse (*green*) gut microbiota using body mass as a proxy of age. Body mass ranges 3.5–33.1g for lab mice and 5.4–28.5g for wild mice. Chronological age is shown in *blue* for the first 30 days for lab mice, in which age–body mass relationship is very clear and not yet strongly influenced by sex differences. Circles in A and C indicate maternal separation status (*hollow* = not yet separated, *filled* = separated). (**A–B**) Quadratic plateau models on asymptotic ASV richness. Vertical lines indicate critical points of plateau. (**A**) Lab mice: *R*^2^=0.596, critical point of inflexion=13.3g. (**B**) Wild mice: *R^2^*=0.049, critical point of inflexion=20.8g. (**C–D)** Quadratic plateau models on the relationship between Aitchison distance to reference juvenile microbiota and body mass. Reference juvenile microbiota was measured separately for laboratory and wild mice by taking the mean of all taxon abundances in mice ≤14 days of age (laboratory mice, *n*=6; body mass ≤6.5g), or ≤7.9g of body mass (wild mice, *n*=16), which is the estimated equivalent of 14 days of age in wild mice. Vertical lines indicate critical points of inflection. (**C**) Lab mice: *R^2^*=0.709, critical point of inflexion=13.0g. (**D**) Wild mice: *R^2^*=0.253, critical point of inflexion=13.2g. (**E**) Beta diversity (based on Jaccard distance) in lab and wild mice. Samples were assigned into classes by dividing the range of body masses across all (lab and wild) samples into three at equal intervals: 3.5–13.3g, 13.4–23.2g, and 23.3–33.1g. Significance was tested with permutational Wilcoxon rank sum test (***; *p*<0.001; *, *p*=0.002).

Body mass was also associated with gut microbial composition in both systems. We investigated gut microbiota maturation by measuring each sample’s compositional distance to a “reference microbiota of a juvenile mouse” (see Methods). Distance to the early life microbiota in each system increased with age, with a plateau reached at around 13g body mass in both systems (corresponding to ∼27 days of age in both lab and wild mice; Fig. S2; Fig. 1F– G). Microbiota variability among individuals decreased with age in both systems, although the pattern showed a more linear decline with increasing body mass in wild mice than lab mice (Fig. 2E).

### Conserved taxon-dependent gut microbiota assembly patterns

Since the major bacterial phyla were conserved across lab and wild mice (Fig. 1C), we compared early life dynamics of the three predominant phyla (Firmicutes, Bacteroidota, and Proteobacteria). These phyla displayed very similar early life trends in both systems, with the relative abundances of Firmicutes and Proteobacteria decreasing and that of Bacteroidota increasing during early life (Fig. 3A–B). The ratio between Firmicutes and Bacteroidota relative abundance became more equal at around 10–15 grams for both lab mice and wild mice. The microbial family Muribaculaceae (previously known as S24–7; unknown genus-level taxonomy) was a key driver of Bacteroidota relative abundance increase in both lab and wild mice (Fig. S4C–D). Declines in the relative abundance of Firmicutes appeared to be driven by a decline in the genus *Ligilactobacillus,* particularly in lab mice (Fig. S4A–B). In addition to *Ligilactobacillus*, taxa from *Limosilactobacillus* and Lachnospiraceae NK4A136 group were key drivers of the reduction in Firmicutes in wild mice (Fig. S4B).

The relative abundance of Proteobacteria decreased in both systems and stabilised at around 10g estimated body mass for lab mice and 15g body mass for wild mice (Fig. 3A–B). In both systems, this reduction in Proteobacteria was driven by taxa belonging to the Enterobacteriaceae family (namely *Escherichia/Shigella*, *Muribacter* and an unknown genus in Enterobacteriaceae; Fig. S5). Finally, shared patterns were also observed in within-phylum ASV richness. In both lab and wild mice, ASVs richness within the phyla Firmicutes and Bacteroidota increased in early life while Proteobacteria ASV richness remained consistently low (Fig. 3C–D). Similar phylum-specific age patterns were also observed when only ASVs that were unique to each system were included (Fig. S6).

**Figure 3.**
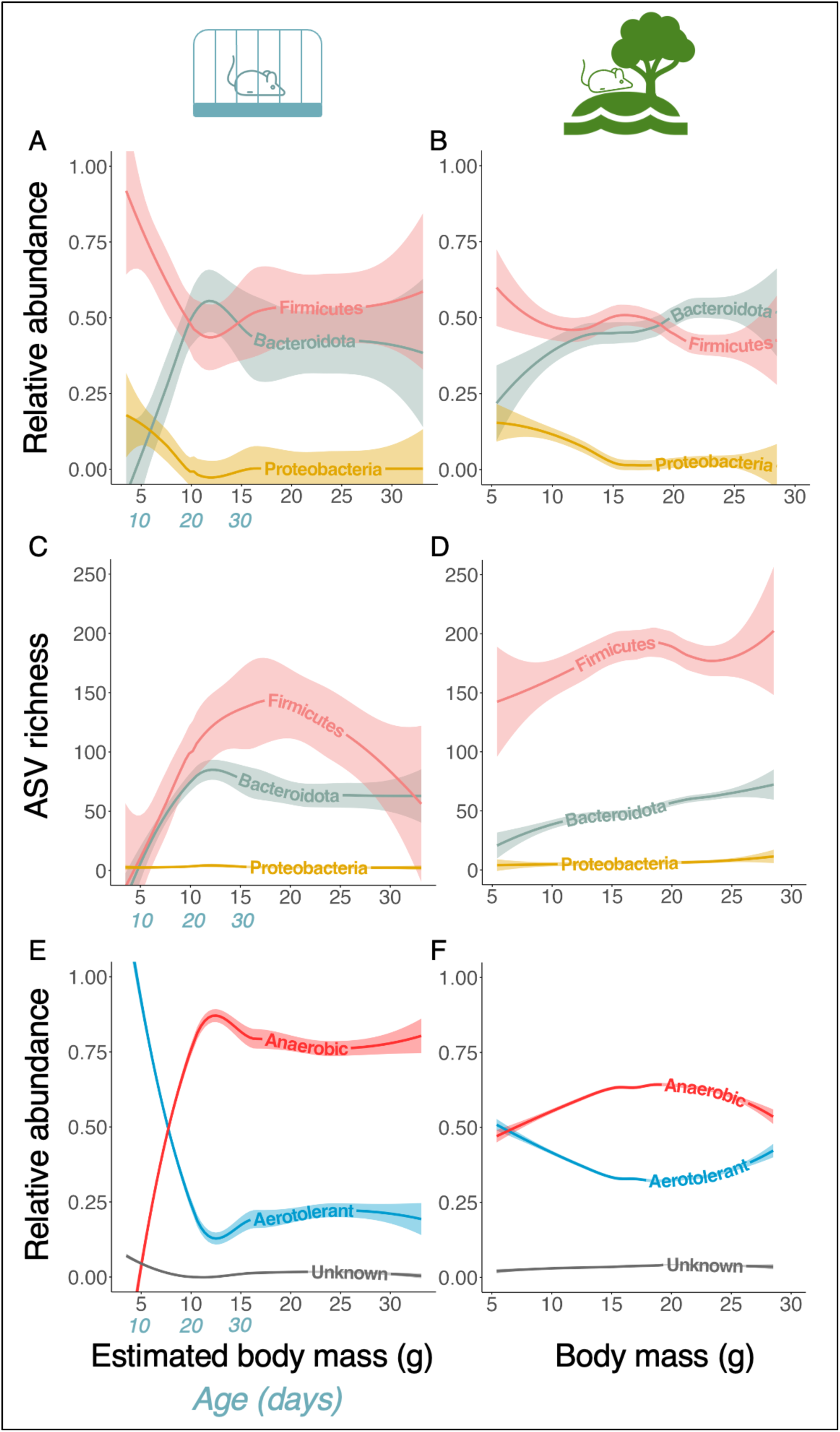
Age-related gut microbial dynamics in (**A**, **C**, **E**) lab and (**B**, **D**, **F**) wild mice using body mass as a proxy of age. Body mass ranges 3.5–33.1g for lab mice and 5.4–28.5g for wild mice. Chronological age is shown in *blue* for the first 30 days for lab mice, in which age–body mass relationship is very clear and not yet strongly influenced by sex differences. (**A–B**) Relative abundance of predominant phyla (only top three phyla with highest total abundance in both systems are presented; abundances were measured from all taxa). (**C–D**) ASV richness (total count of unique ASVs) in predominant phyla. Lines are locally estimated scatterplot smoothing (LOESS) lines with 95% confidence interval bands. (**E–F**) Relative abundance of aerotolerant and obligate anaerobic bacteria across body mass in (**E**) lab and (**F**) wild mice respectively. Lines are locally estimated scatterplot smoothing (LOESS) lines with 95% confidence interval bands. *Anaerobic* = obligate anaerobes, *aerotolerant* = everything else with known aerotolerance, *unknown* = bacteria with unknown aerotolerance.

### Age-related changes in gut microbial phenotypes and functional pathways

As the mammalian gut becomes increasingly anoxic in early life^53^, we hypothesised that the ratio between aerotolerant and anaerobic bacteria would shift in early life, favouring anaerobes. In lab mice, we detected a clear age-related pattern in bacterial aerotolerance consistent with this expectation. Aerotolerant bacteria initially dominated the gut microbiota but declined in relative abundance while anaerobes increased, eventually dominating over aerotolerant taxa in relative abundance by a ratio of approximately 3:1 by ∼15g body mass (Fig. 3E). In wild mice, we similarly observed an increase in anaerobes and a decrease in aerotolerant taxa, although the pattern was less pronounced. Further, aerotolerant taxa retained a higher relative abundance in wild compare to lab mice in adulthood (mean relative abundance of aerotolerant taxa in mice >20g: 34.2% in wild vs 23.1% in lab; Fig. 3F).

Finally, we investigated predicted functional profiles of the gut microbiota. The relative abundance of functional pathways showed a clear shift at around 8g body mass (17d of age) in lab mice (Fig. 4C). This therefore occurred 2–5 days before pups were separated from their mother and coincided with a clear compositional shift in the bacterial genera present (Fig. 4A). No clear shift in functional profile or bacterial genus composition was detected in wild mice (Fig. 4B, 4D). Given this finding of a clear compositional shift in lab mice during the weaning period, we re-assessed trends in phylum composition and aerotolerance in lab mice according to whether they had been separated from dams and thus were eating an entirely solid food-based diet. This indicated that much of the observed age-related gut microbial changes we observed occurred before pups were separated from dams, and therefore fully weaned (Fig. S7).

**Figure 4.**
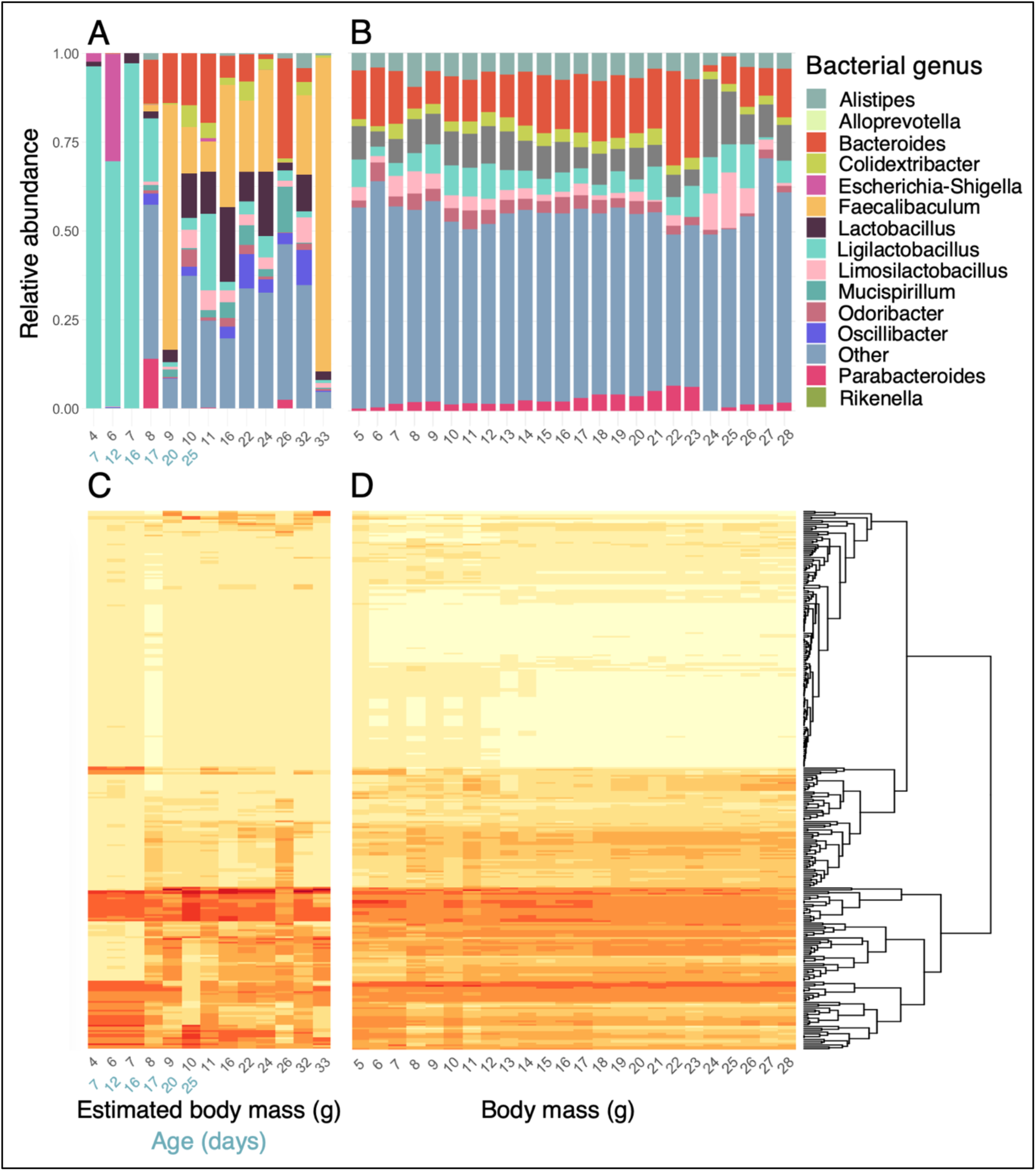
Composition of bacterial genera and functional pathways in laboratory and wild mice. Body mass (used as a proxy of age) ranges 3.5–33.1g for lab mice and 5.4–28.5g for wild mice. Chronological age is shown in *blue* for the first 30 days for lab mice, in which age–body mass relationship is very clear and not yet strongly influenced by sex differences. **(A–B)** Mean relative abundance of bacterial genera across each available unit of (estimated) body mass in (**A**) lab and (**B**) wild mice. (**C–D**) Mean relative abundance of predicted functional pathways across each available unit of age/body mass in (**C**) lab and (**D**) wild mice. Rows are unique MetaCyc pathways ordered by hierarchical clustering as indicated with dendrogram. Colour darkness increases with relative abundance (range 0.0–2.7%).

## Discussion

Considering laboratory mice are widely used in research on the gut microbiota and its links to host physiology, we set out to investigate how well the dynamics of gut microbiota maturation in lab mice represent those occurring in wild members of the same species. Several key patterns in microbial community assembly were conserved across settings, despite wild mice having higher microbiota richness and distinct taxonomic composition, as reported in previous studies comparing adult lab and wild mice^54–57^.

Our results illustrate how in both the lab and the wild, the gut microbiota of house mice matured during the first month of life. During this time, the microbiota increased in alpha diversity, changed in taxonomic composition, and became more homogenous in composition among individuals. These patterns resemble those of gut microbiota maturation reported for humans and wild gelada baboons^20,25,27,58,59^ but contrast those reported in a study of wild chimpanzees, where richness declined during early life^26^. As previously documented in humans^20,60^, anaerobic bacteria also increased in relative abundance during early life, likely because they are best adapted to the increasingly anoxic conditions of the developing gut.

Despite having taxonomically distinct gut microbiotas at nearly all levels examined, phylum-level composition remained similar among lab and wild mice, as reported by previous studies^54,56^. We show that these phyla also show remarkably consistent age-related changes in relative abundance and richness, irrespective of whether mice were lab-housed or wild. Moreover, while shared taxa (ASVs detected in both lab and wild mice) made up nearly half the gut microbiota in both settings, age-related patterns in phylum relative abundances and richness were unchanged when only taxa unique to each study system were analysed. These consistent taxonomic shifts are thus not solely driven by a set of shared ASVs, but appear to reflect general trends occurring in members of these phyla. In both systems, the ratio between Firmicutes and Bacteroidota decreased as young mice matured, coinciding with an increase in the ratio of anaerobic to aerotolerant bacteria. The decrease in Firmicutes relative abundance with age was primarily driven by declining *Ligilactobacillus* in lab mice and *Ligilactobacillus* and *Limosilactobacillus* in wild mice. These aerotolerant taxa have been detected in the milk of various mammal species including mice^61–64^ and may be transmitted by lactation, such that their reduction may be driven by declining milk consumption in weaning mice^65^. *Ligilactobacillus* had remarkably low relative abundance in wild mice of all ages, which would be expected if this taxon was primarily transmitted through lactation, as wild mice must have been either weaning or fully weaned to be trapped in this study. The age-related increase in Bacteroidota relative abundance was largely driven by Muribaculaceae in both systems. These are anaerobic fermentative bacteria with a broad repertoire of carbohydrate metabolising capabilities^66^ that appear to be socially and vertically transmitted in wild wood mice^67,68^. While we can only speculate on the mechanisms underlying this increase, two possibilities are increased consumption of a plant-based diet and increasing social interaction as young wild mice mature.

Changes in the relative abundance and richness of proteobacterial taxa were also very similar in lab and wild mice. Consistent with documented patterns in humans, primates, and mice^20,24,28,69^, the relative abundance of Proteobacteria decreased with age, with Enterobacteriaeae playing a prominent role in this trend. The precise drivers of this pattern remain to be explored, but could include declining oxygen levels in the gut (as Enterobacteriacaeae are facultative aerobes^70^), or immune maturation, as this family harbours several genera with pathogenic potential, and early life immune maturation should increase the host’s ability to suppress such pathogenic taxa. Indeed, *Escherichia,* a genus containing well-known pathogenic species, was a key driver of the early-life decline in Enterobacteriacaeae among lab mice.

While several broad patterns of alpha and beta diversity change during early life were conserved among lab and wild mice, microbiota maturation appeared to be more gradual among wild mice. In lab mice, alpha diversity reached a plateau rather abruptly at the same time compositional maturation was completed at around 27 days of age. In wild mice, however, alpha diversity appeared to increase up to a body mass of 20g, at which point wild mice are expected to be ∼40–80 days old. Several factors may underlie the more gradual patterns of gut microbiota maturation observed in wild mice. One possibility is that weaning (gradual transition from a purely milk-based to a purely solid food diet), a crucial process driving gut microbiota maturation in mammals^20,24,71^, was captured more completely in our lab compared to wild mouse dataset. Laboratory mice have been shown to start nibbing solid food from 12 days of age^72^ and have been shown to wean between days 17 and 22 of age^73^. While alpha and beta diversity in the lab plateaued a few days after pups were separated from their mothers, shifts in microbiota composition and diversity had already begun before the expected start of solid food consumption and several aspects of maturation were completed before maternal separation. In particular, the lab mouse gut microbiota displayed a clear shift in predicted microbiota functions at 17d of age when weaning is expected to have ramped up, which coincided with a marked shift in the microbial genera present, including a stark decrease in milk-associated *Ligilactobacillus*. Together these patterns support previous findings that the introduction of solid food or cessation of milk consumption alone can induce certain changes^33,20,24,71^, but also suggest that the overall maturation of the gut microbiota is driven by the gradual transition from a milk-based diet to a solid food diet.

In our study, the lab dataset included mice still on a purely milk-based diet, while wild mice included were independent enough to be trapped and thus expected to be either fully weaned or in the late stages of weaning. As such, we may have captured only the tail-end of gut microbial maturation in wild mice, which could contribute to the maturation appearing more gradual in the wild, and explain why an age-related functional shift was only detected in lab mice. Further studies where wild/semi-wild pups can be sampled in nests, allowing capture of the full successional process, would be useful to investigate whether and when a similar functional shift occurs in more natural settings, and whether gut microbiota assembly is indeed more gradual in the wild.

Another factor that could contribute to more gradual trends of microbiota maturation among wild mice is differing patterns of microbial exposure. Highly controlled lab conditions (including confined space, pathogen exclusion, a small number of cage mates, and a homogeneous chow diet) should lead to a restricted pool of microbes available for colonisation, which could generate a faster saturation of alpha diversity as the available microbial pool is explored. In contrast, young wild mice should interact with a much more dynamic, variable microbial landscape as they disperse, consume a variable diet, and establish their adult home ranges and social relationships. This could extend their exposure to a variety of environmental and conspecific microbial reservoirs, potentially contributing to a more gradual rise in alpha diversity during early life.

Another notable difference in the early life gut microbiota of lab and wild mice involved aerotolerance phenotypes of the constituent bacteria. In both settings the relative abundance of anaerobes increased while that of aerotolerant bacteria declined with age. This pattern was again much more striking in lab mice than wild mice, while in wild mice anaerobes had higher relative abundance than aerotolerant taxa after brief dominance of aerotolerant taxa in very early life. This may again be largely due to a lack of unweaned mice in our wild dataset, such that the earliest part of microbiota maturation (when the gut is dominated by aerotolerant taxa) was not unobserved. Once maturation was complete, however, lab and wild mice showed a clear difference in the ratio of aerotolerant to anaerobic microbes, with the relative abundance of aerotolerant gut microbes remaining much higher in wild than lab mice. Several factors may underlie this difference. The abundant aerotolerant taxa in wild mice were not obviously pathogenic species, primarily comprising of members of the families Bacteroidaceae and Lactobacillaceae. As such, increased pathogen exposure in the wild does not appear to explain this difference. However, differences in microbial exposures (if wild mice are exposed to a greater number of aerotolerant and/or sporulating microbial taxa that persist in their complex natural environment) or diet differences (if lab mouse chow favours a more anaerobic fermentative environment than would occur in wild mice) could both play a role. Further work to characterise the diet of wild mice, and microbes in their abiotic environment could shed further light on this interesting difference.

Together, our finding that several major trends in gut microbiota assembly are conserved across lab and wild mice indicates that to a large extent, the broad patterns of gut microbiota maturation follow an intrinsic host programme that transcends contrasting genetic and environmental backgrounds, and the existence of different component species. Our results therefore indicate that meaningful insights can be drawn from lab models to understand the developing gut microbiota’s significance in early life. Some apparent differences between lab and wild gut microbiota maturation warrant further attention, however, such as the potentially more gradual changes in wild animals and greater abundance of aerotolerant taxa into adulthood. Further understanding these differences and the drivers of gut microbiota assembly patterns in wild mice could allow adjustment of lab mouse husbandry to achieve assembly of gut microbiota that more closely resembles that occurring in natural populations, allowing more accurate inference about the impact of this key microbial developmental stage on mammalian biology.

## Supporting information

References for Supplementary Table 1

## Acknowledgements

We thank Giselle Eagle and Richard Brown (Wardens of Skokholm Island), the Friends of Skokholm and Skomer, the Wildlife Trust of South and West Wales and field assistants Billy Dykes and Olivia Pargeter for their help in enabling the Skokholm wild mouse data collection.

## Author information

EH and SCLK set up the wild mouse study system. EH, AR and SCLK collected the data. EH conducted the laboratory work and analysed the data. EH wrote the manuscript. All authors contributed to the final manuscript.

## Ethical statement

Wild mouse data collection was done under Home Office licence PPL PB0178858 held at the University of Oxford, and with a research permit from the Islands Conservation Advisory Committee (ICAC), and Natural Resources Wales. Mice were not subject to intervention as part of this study.

## Data and code availability

The 16S rRNA amplicon sequencing data used in this study have been deposited in GenBank under SRA accession: PRJNA1028479. Code used in the study will be made publicly available upon publication.

## References

1. Visconti A, Le Roy CI, Rosa F, et al. Interplay between the human gut microbiome and host metabolism. Nat Commun. 2019;10(1):4505. doi:10.1038/s41467-019-12476-z

2. Turnbaugh PJ, Ley RE, Mahowald MA, Magrini V, Mardis ER, Gordon JI. An obesity-associated gut microbiome with increased capacity for energy harvest. Nature. 2006;444(7122):1027–1031. doi:10.1038/nature05414

3. Liberti J, Kay T, Quinn A, et al. The gut microbiota affects the social network of honeybees. Nat Ecol Evol. 2022;6(10):1471–1479. doi:10.1038/s41559-022-01840-w

4. Rudolph K, Schneider D, Fichtel C, Daniel R, Heistermann M, Kappeler PM. Drivers of gut microbiome variation within and between groups of a wild Malagasy primate. Microbiome. 2022;10(1):28. doi:10.1186/s40168-021-01223-6

5. Nishida AH, Ochman H. Rates of gut microbiome divergence in mammals. Mol Ecol. 2018;27(8):1884–1897. doi:10.1111/mec.14473

6. Vandeputte D, Kathagen G, D’hoe K, et al. Quantitative microbiome profiling links gut community variation to microbial load. Nature. 2017;551(7681):507–511. doi:10.1038/nature24460

7. Stewart CJ, Ajami NJ, O’Brien JL, et al. Temporal development of the gut microbiome in early childhood from the TEDDY study. Nature. 2018;562(7728):583–588. doi:10.1038/s41586-018-0617-x

8. Yassour M, Vatanen T, Siljander H, et al. Natural history of the infant gut microbiome and impact of antibiotic treatment on bacterial strain diversity and stability. Sci Transl Med. 2016;8(343):343ra81. doi:10.1126/scitranslmed.aad0917

9. Depner M, Taft DH, Kirjavainen P V., et al. Maturation of the gut microbiome during the first year of life contributes to the protective farm effect on childhood asthma. Nat Med. 2020;26(11):1766–1775. doi:10.1038/s41591-020-1095-x

10. Gonze D, Coyte KZ, Lahti L, Faust K. Microbial communities as dynamical systems. Curr Opin Microbiol. 2018;44:41–49. doi:10.1016/j.mib.2018.07.004

11. Chung H, Pamp SJ, Hill JA, et al. Gut immune maturation depends on colonization with a host-specific microbiota. Cell. 2012;149(7):1578–1593. doi:10.1016/j.cell.2012.04.037

12. Sharon G, Sampson TR, Geschwind DH, Mazmanian SK. The Central Nervous System and the Gut Microbiome. Cell. 2016;167(4):915–932. doi:10.1016/j.cell.2016.10.027

13. Olin A, Henckel E, Chen Y, et al. Stereotypic Immune System Development in Newborn Children. Cell. 2018;174(5):1277–1292.e14. doi:10.1016/j.cell.2018.06.045

14. Darabi B, Rahmati S, HafeziAhmadi MR, Badfar G, Azami M. The association between caesarean section and childhood asthma: an updated systematic review and meta-analysis. *Allergy*, Asthma & Clinical Immunology. 2019;15(1):62. doi:10.1186/s13223-019-0367-9

15. Bäckhed F, Roswall J, Peng Y, et al. Dynamics and Stabilization of the Human Gut Microbiome during the First Year of Life. Cell Host Microbe. 2015;17(5):690–703. doi:10.1016/j.chom.2015.04.004

16. Ferretti P, Pasolli E, Tett A, et al. Mother-to-Infant Microbial Transmission from Different Body Sites Shapes the Developing Infant Gut Microbiome. Cell Host Microbe. 2018;24(1):133–145.e5. doi:10.1016/j.chom.2018.06.005

17. Olin A, Henckel E, Chen Y, et al. Stereotypic Immune System Development in Newborn Children. Cell. 2018;174(5):1277–1292.e14. doi:10.1016/j.cell.2018.06.045

18. Lubin JB, Green J, Maddux S, Brodsky IE, Planet PJ, Silverman MA. Arresting microbiome development limits immune system maturation and resistance to infection in mice. Cell Host Microbe. 2023;31:554–570.e7. doi:10.1016/j.chom.2023.03.006

19. Shao Y, Forster SC, Tsaliki E, et al. Stunted microbiota and opportunistic pathogen colonization in caesarean-section birth. Nature. 2019;574(7776):117–121. doi:10.1038/s41586-019-1560-1

20. Bäckhed F, Roswall J, Peng Y, et al. Dynamics and Stabilization of the Human Gut Microbiome during the First Year of Life. Cell Host Microbe. 2015;17(5):690–703. doi:10.1016/j.chom.2015.04.004

21. Lehtimäki J, Karkman A, Laatikainen T, et al. Patterns in the skin microbiota differ in children and teenagers between rural and urban environments. Sci Rep. 2017;7(1):45651. doi:10.1038/srep45651

22. Ruokolainen L, Paalanen L, Karkman A, et al. Significant disparities in allergy prevalence and microbiota between the young people in Finnish and Russian Karelia. Clin Exp Allergy. 2017;47(5):665–674. doi:10.1111/cea.12895

23. Ruokolainen L, Parkkola A, Karkman A, et al. Contrasting microbiotas between Finnish and Estonian infants: Exposure to Acinetobacter may contribute to the allergy gap. Allergy. 2020;75(9):2342–2351. doi:10.1111/all.14250

24. Baniel A, Petrullo L, Mercer A, et al. Maternal effects on early-life gut microbiota maturation in a wild nonhuman primate. Curr Biol. 2022;32(20):4508–4520.e6. doi:10.1016/j.cub.2022.08.037

25. Petrullo L, Baniel A, Jorgensen MJ, Sams S, Snyder-Mackler N, Lu A. The early life microbiota mediates maternal effects on offspring growth in a nonhuman primate. iScience. 2022;25(3):103948. doi:10.1016/j.isci.2022.103948

26. Reese AT, Phillips SR, Owens LA, et al. Age Patterning in Wild Chimpanzee Gut Microbiota Diversity Reveals Differences from Humans in Early Life. Current Biology. 2021;31(3):613–620.e3. doi:10.1016/J.CUB.2020.10.075

27. Pantoja-Feliciano IG, Clemente JC, Costello EK, et al. Biphasic assembly of the murine intestinal microbiota during early development. ISME J. 2013;7(6):1112–1115. doi:10.1038/ismej.2013.15

28. Li X, Ren Y, Zhang J, et al. Development of Early-Life Gastrointestinal Microbiota in the Presence of Antibiotics Alters the Severity of Acute DSS-Induced Colitis in Mice. Yeruva L, ed. Microbiol Spectr. 2022;10(3). doi:10.1128/spectrum.02692-21

29. Martínez I, Maldonado-Gomez MX, Gomes-Neto JC, et al. Experimental evaluation of the importance of colonization history in early-life gut microbiota assembly. Elife. 2018;7. doi:10.7554/eLife.36521

30. Tochitani S, Ikeno T, Ito T, Sakurai A, Yamauchi T, Matsuzaki H. Administration of Non-Absorbable Antibiotics to Pregnant Mice to Perturb the Maternal Gut Microbiota Is Associated with Alterations in Offspring Behavior. Shankar K, ed. PLoS One. 2016;11(1):e0138293. doi:10.1371/journal.pone.0138293

31. Laukens D, Brinkman BM, Raes J, De Vos M, Vandenabeele P. Heterogeneity of the gut microbiome in mice: guidelines for optimizing experimental design. Normark BH, ed. FEMS Microbiol Rev. 2016;40(1):117–132. doi:10.1093/femsre/fuv036

32. Schloss PD, Schubert AM, Zackular JP, Iverson KD, Young VB, Petrosino JF. Stabilization of the murine gut microbiome following weaning. Gut Microbes. 2012;3(4):383–393. doi:10.4161/gmic.21008

33. Russell AL, McAdams ZL, Donovan E, et al. The contribution of maternal oral, vaginal, and gut microbiota to the developing offspring gut. Sci Rep. 2023;13(1):13660. doi:10.1038/s41598-023-40703-7

34. Hughes KR, Schofield Z, Dalby MJ, et al. The early life microbiota protects neonatal mice from pathological small intestinal epithelial cell shedding. The FASEB Journal. 2020;34(5):7075–7088. doi:10.1096/fj.202000042R

35. Berry RJ, Jakobson ME. Life and Death in an Island Population of the House Mouse. 1971;(June 1967):187–197.

36. Berry RJ, Lidicker W. Ageing in an island population of the House mouse. 1973;(August). doi:10.1093/ageing/2.4.235

37. Spangenberg E, Wallenbeck A, Eklöf AC, Carlstedt-Duke J, Tjäder S. Housing breeding mice in three different IVC systems: maternal performance and pup development. Lab Anim. 2014;48(3):193–206. doi:10.1177/0023677214531569

38. Body Weight Information for C57BL/6J | The Jackson Laboratory. Accessed August 17, 2023. https://www.jax.org/jax-mice-and-services/strain-data-sheet-pages/body-weight-chart-000664

39. Gray MM, Parmenter MD, Hogan CA, et al. Genetics of Rapid and Extreme Size Evolution in Island Mice. Genetics. 2015;201(1):213–228. doi:10.1534/genetics.115.177790

40. Ferrari M, Lindholm AK, König B. The risk of exploitation during communal nursing in house mice, Mus musculus domesticus. Anim Behav. 2015;110:133–143. doi:10.1016/J.ANBEHAV.2015.09.018

41. Parada AE, Needham DM, Fuhrman JA. Every base matters: assessing small subunit rRNA primers for marine microbiomes with mock communities, time series and global field samples. Environ Microbiol. 2016;18(5):1403–1414. doi:10.1111/1462-2920.13023

42. Walters W, Hyde ER, Berg-Lyons D, et al. Improved Bacterial 16S rRNA Gene (V4 and V4-5) and Fungal Internal Transcribed Spacer Marker Gene Primers for Microbial Community Surveys. mSystems. 2016;1(1). doi:10.1128/mSystems.00009-15

43. Comeau AM, Douglas GM, Langille MGI. Microbiome Helper: a Custom and Streamlined Workflow for Microbiome Research. Eisen J, ed. mSystems. 2017;2(1). doi:10.1128/mSystems.00127-16

44. Callahan BJ, McMurdie PJ, Rosen MJ, Han AW, Johnson AJA, Holmes SP. DADA2: High-resolution sample inference from Illumina amplicon data. Nat Methods. 2016;13(7):581–583. doi:10.1038/nmeth.3869

45. Hsieh TC MKCA. iNEXT: Interpolation and Extrapolation for Species Diversity. R package version 3.0.0. Published online 2022.

46. McMurdie PJ, Holmes S. phyloseq: an R package for reproducible interactive analysis and graphics of microbiome census data. PLoS One. 2013;8(4):e61217. doi:10.1371/journal.pone.0061217

47. Davis NM, Proctor DM, Holmes SP, Relman DA, Callahan BJ. Simple statistical identification and removal of contaminant sequences in marker-gene and metagenomics data. Microbiome. 2018;6(1):226. doi:10.1186/s40168-018-0605-2

48. Douglas GM, Maffei VJ, Zaneveld JR, et al. PICRUSt2 for prediction of metagenome functions. Nat Biotechnol. 2020;38(6):685–688. doi:10.1038/s41587-020-0548-6

49. Oksanen, F.J., Simpson G.L., Blanchet GF. et al. vegan: Community Ecology Package [R package vegan version 2.6-4]. Published online October 11, 2017. https://cran.r-project.org/web/packages/vegan/index.html

50. Lahti L, Shetty S. Tools for microbiota analysis in R. Published online 2017.

51. Maintainer EA, Arnhold E. Package “easynls” Type Package Title Easy Nonlinear Model.; 2022. Accessed August 17, 2023. https://cran.r-project.org/web/packages/easynls/easynls.pdf

52. Wickham H. ggplot2: Elegant Graphics for Data Analysis. Published online 2016.

53. Albenberg L, Esipova T V, Judge CP, et al. Correlation between intraluminal oxygen gradient and radial partitioning of intestinal microbiota. Gastroenterology. 2014;147(5):1055–63.e8. doi:10.1053/j.gastro.2014.07.020

54. Rosshart SP, Herz J, Vassallo BG, et al. Laboratory mice born to wild mice have natural microbiota and model human immune responses. Science (1979). 2019;365(6452). doi:10.1126/science.aaw4361

55. Wang J, Linnenbrink M, Künzel S, et al. Dietary history contributes to enterotype-like clustering and functional metagenomic content in the intestinal microbiome of wild mice. Proc Natl Acad Sci U S A. 2014;111(26):E2703–10. doi:10.1073/pnas.1402342111

56. Kreisinger J, Cížková D, Vohánka J, Piálek J. Gastrointestinal microbiota of wild and inbred individuals of two house mouse subspecies assessed using high-throughput parallel pyrosequencing. Mol Ecol. 2014;23(20):5048–5060. doi:10.1111/mec.12909

57. Wang J, Kalyan S, Steck N, et al. Analysis of intestinal microbiota in hybrid house mice reveals evolutionary divergence in a vertebrate hologenome. Nat Commun. 2015;6(1):6440. doi:10.1038/ncomms7440

58. Derrien M, Alvarez AS, de Vos WM. The Gut Microbiota in the First Decade of Life. Trends Microbiol. 2019;27(12):997–1010. doi:10.1016/j.tim.2019.08.001

59. Yatsunenko T, Rey FE, Manary MJ, et al. Human gut microbiome viewed across age and geography. Nature. 2012;486(7402):222–227. doi:10.1038/nature11053

60. Guittar J, Shade A, Litchman E. Trait-based community assembly and succession of the infant gut microbiome. Nat Commun. 2019;10(1):512. doi:10.1038/s41467-019-08377-w

61. Quilodrán-Vega S, Albarracin L, Mansilla F, et al. Functional and Genomic Characterization of Ligilactobacillus salivarius TUCO-L2 Isolated From Lama glama Milk: A Promising Immunobiotic Strain to Combat Infections. Front Microbiol. 2020;11:608752. doi:10.3389/fmicb.2020.608752

62. Qi C, Tu H, Zhao Y, et al. Breast Milk-Derived *Limosilactobacillus reuteri* Prevents Atopic Dermatitis in Mice via Activating Retinol Absorption and Metabolism in Peyer’s Patches. Mol Nutr Food Res. 2023;67(2):2200444. doi:10.1002/mnfr.202200444

63. Stinson LF, Sindi ASM, Cheema AS, et al. The human milk microbiome: who, what, when, where, why, and how? Nutr Rev. 2021;79(5):529–543. doi:10.1093/nutrit/nuaa029

64. de Andrés J, Jiménez E, Chico-Calero I, Fresno M, Fernández L, Rodríguez JM. Physiological Translocation of Lactic Acid Bacteria during Pregnancy Contributes to the Composition of the Milk Microbiota in Mice. Nutrients. 2017;10(1). doi:10.3390/nu10010014

65. Qi C, Zhou J, Tu H, et al. Lactation-dependent vertical transmission of natural probiotics from the mother to the infant gut through breast milk. Food Funct. 2022;13(1):304–315. doi:10.1039/D1FO03131G

66. Lagkouvardos I, Lesker TR, Hitch TCA, et al. Sequence and cultivation study of Muribaculaceae reveals novel species, host preference, and functional potential of this yet undescribed family. Microbiome. 2019;7(1):28. doi:10.1186/s40168-019-0637-2

67. Wanelik KM, Raulo A, Troitsky T, Husby A, Knowles SCL. Maternal transmission gives way to social transmission during gut microbiota assembly in wild mice. Anim Microbiome. 2023;5(1):29. doi:10.1186/s42523-023-00247-7

68. Raulo A, Allen BE, Troitsky T, et al. Social networks strongly predict the gut microbiota of wild mice. ISME J. 2021;15(9):2601–2613. doi:10.1038/s41396-021-00949-3

69. Jokela R, Korpela K, Jian C, et al. Quantitative insights into effects of intrapartum antibiotics and birth mode on infant gut microbiota in relation to well-being during the first year of life. Gut Microbes. 2022;14(1):2095775. doi:10.1080/19490976.2022.2095775

70. Enav H, Bäckhed F, Ley RE. The developing infant gut microbiome: A strain-level view. Cell Host Microbe. 2022;30(5):627–638. doi:10.1016/j.chom.2022.04.009

71. Bergström A, Skov TH, Bahl MI, et al. Establishment of intestinal microbiota during early life: a longitudinal, explorative study of a large cohort of Danish infants. Appl Environ Microbiol. 2014;80(9):2889–2900. doi:10.1128/AEM.00342-14

72. JAX Mice Pup Appearance By Age | The Jackson Laboratory. Accessed August 17, 2023. https://www.huanglabmcgill.org/uploads/1/2/1/6/121601138/lt0001_mouse_pup_appearance.pdf

73. K□nig B, Markl H. Maternal care in house mice. Behav Ecol Sociobiol. 1987;20(1):1–9. doi:10.1007/BF00292161

