## Supplementary Material for "Early-life gut microbiota assembly patterns are conserved between laboratory and wild mice"

### Supplementary data

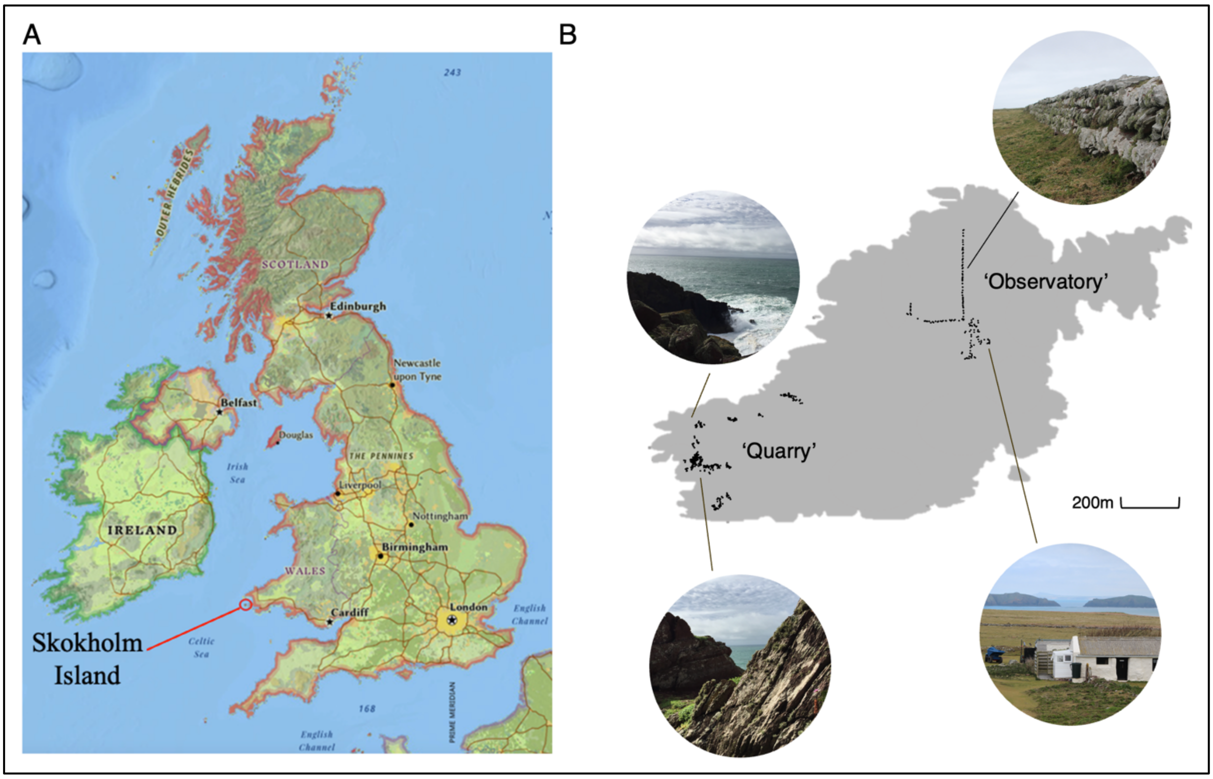


**Supplementary Figure 1.** (**A**) Skokholm Island is located 4 km off the coast of Pembrokeshire in south-west Wales, UK. (**B**) Two wild house mouse sampling sites, 'Observatory' and 'Quarry', on Skokholm Island. 150 trapping points (black circles) were distributed at each site.

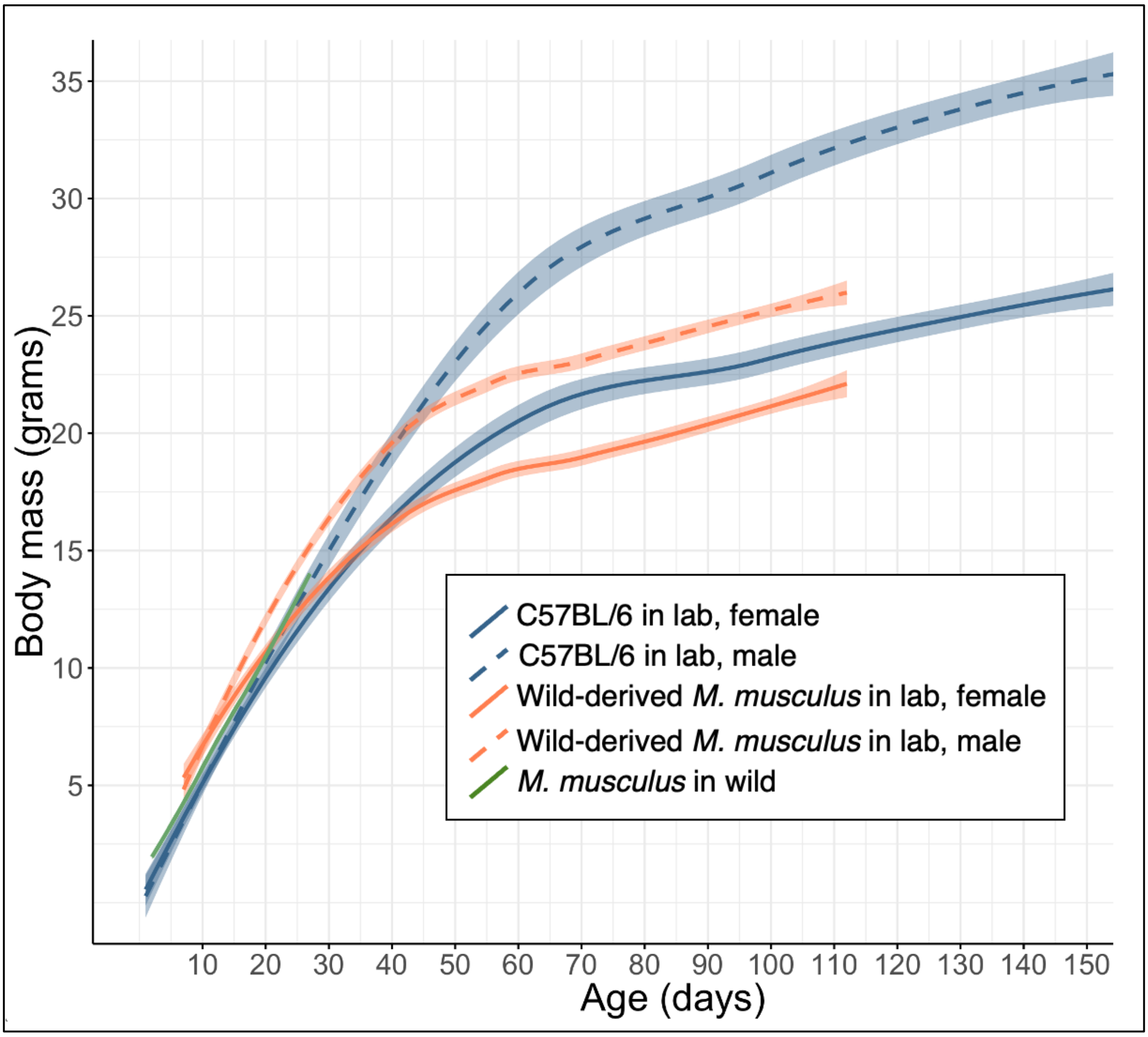


**Supplementary Figure 2.** Age–body mass relationship in house mouse. *Blue:* C57BL/6 (*Mus musculus*) data has been reproduced from Figure 3 in Spangenberg et al (2014)[1] and The Jackson Laboratory website[2]. *Orange:* Wild-derived *M. musculus* (Gough Island, home to the largest wild house mice recorded; mice born in laboratory) data has been reproduced from Figure 3 in Gray et al (2015)[3]. *Green:* Wild *M. musculus* data has been reproduced from Figure 5 in Ferrari et al (2015)[4], and depicts the body masses of young wild mice of known age inhabiting a barn near Zurich, Switzerland. Line type indicates sex: *solid* = female, *dashed* = male.

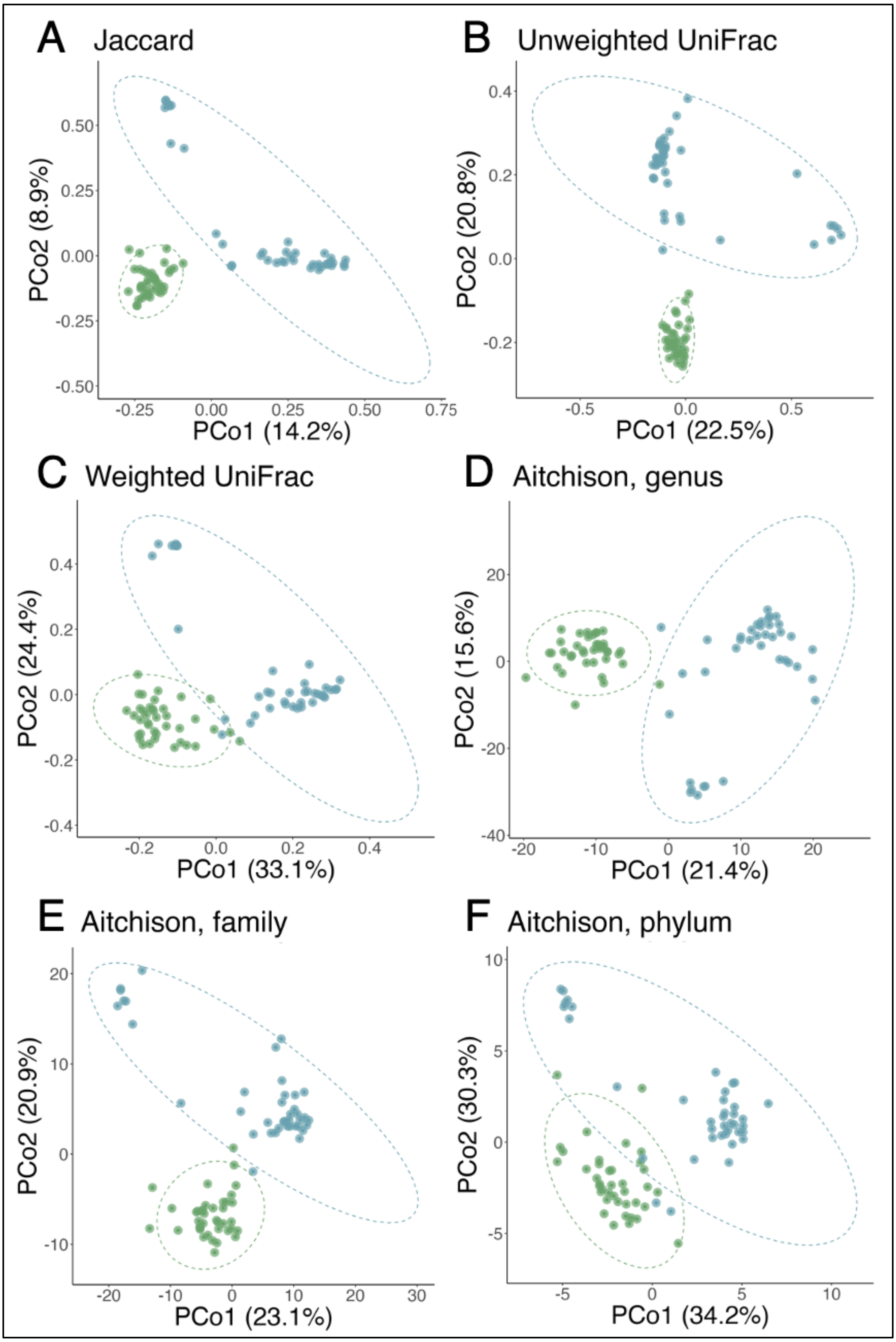


**Supplementary Figure 3.** Principal coordinates analysis (PCoA) of 78 faecal samples from 39 laboratory and 39 wild mice using (**A**) Jaccard, (**B**) unweighted UniFrac, (**C**) weighted and (**D**–**F**) Aitchison distances at (**A–C**) amplicon sequence variant (ASV), (**D**) genus, (**E**) family, or (**F**) phylum level. Colour indicates sample source (*blue* = laboratory, *green* = wild).

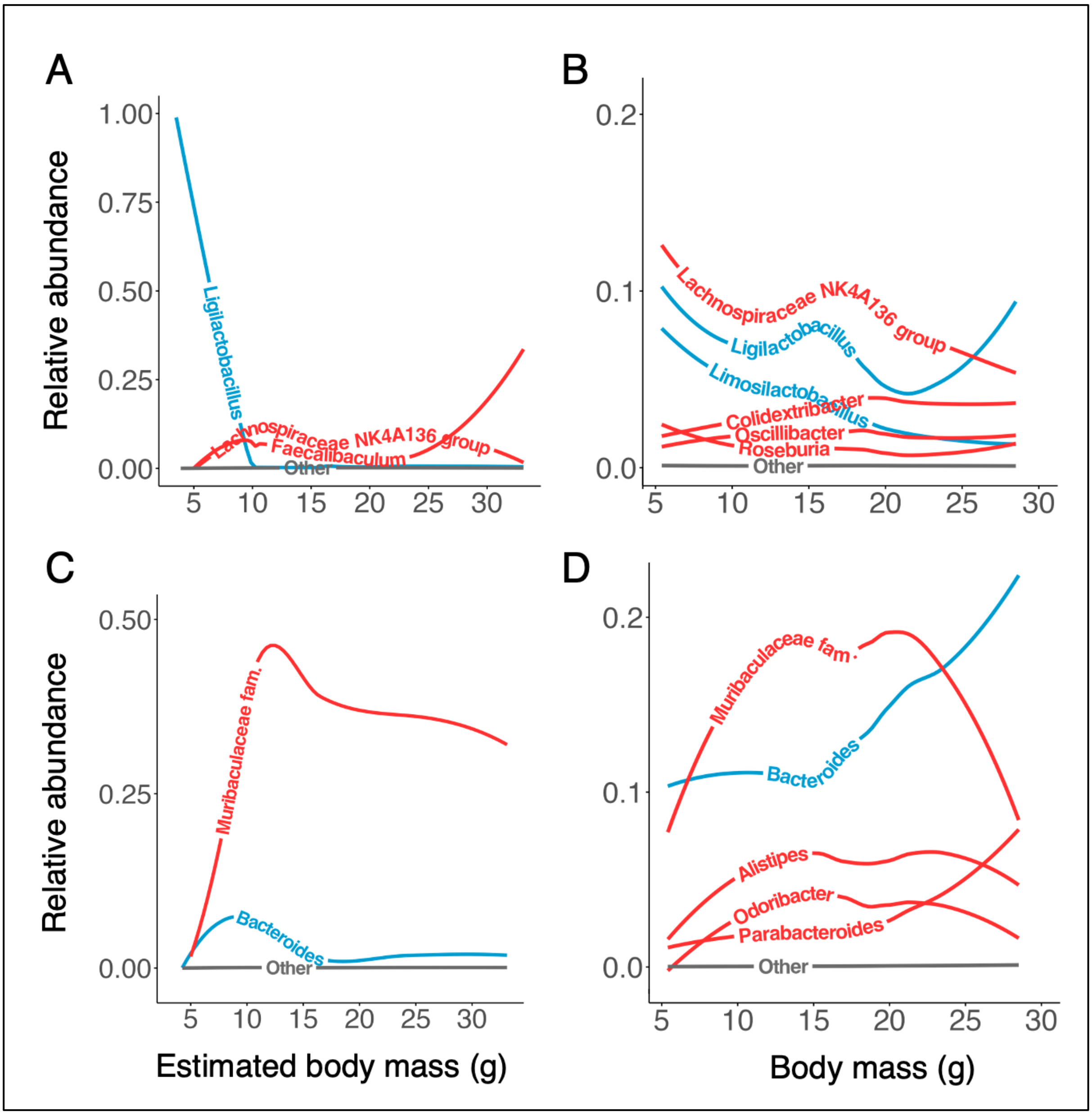


**Supplementary Figure 4.** Relative abundance of genera from (**A–B**) Firmicutes and (**C–D**) Bacteroidota, in (**A, C**) laboratory and (**B, D**) wild mice. Relative abundances are measured from the whole microbiota, rather than within phylum. Lines are locally estimated scatterplot smoothing (LOESS) lines. Confidence intervals are not plotted for easier interpretation. Lines are coloured by genus aerotolerance (*red* = obligate anaerobes, *blue* = aerotolerant, *grey* = unknown aerotolerance). Note the variable y-axes scales to aid visibility of taxa with low abundance.

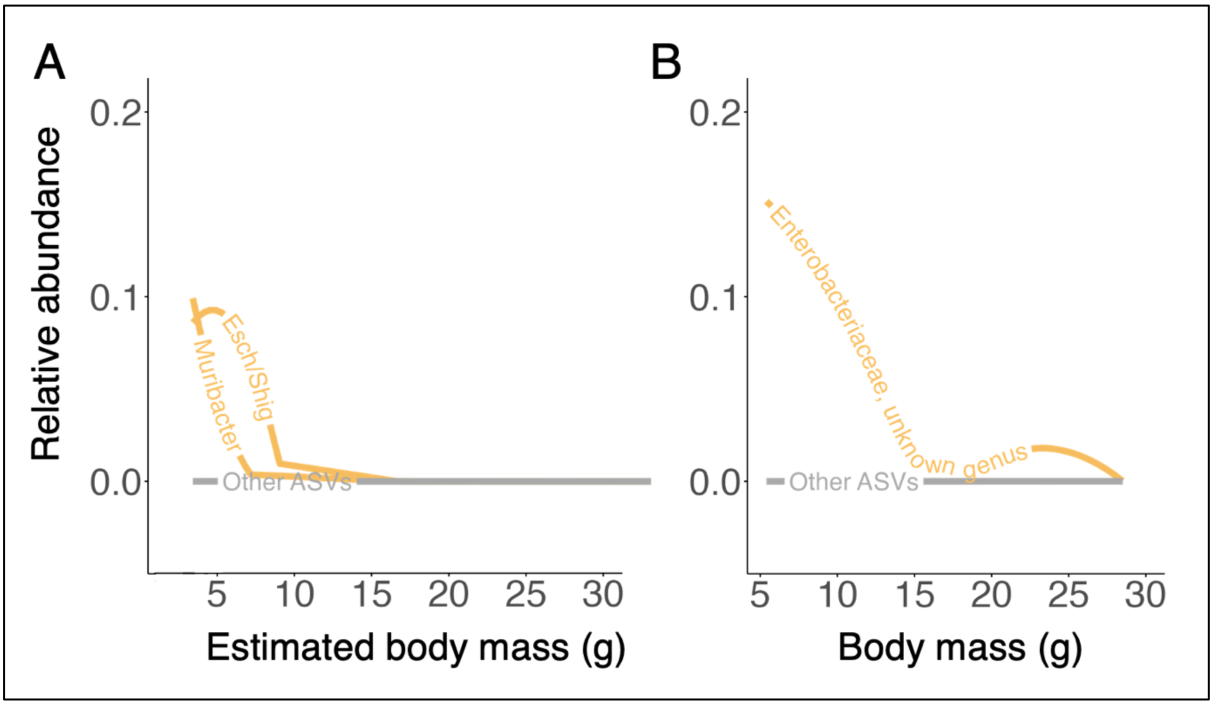


**Supplementary Figure 5.** Relative abundance of ASVs assigned to the phylum Proteobacteria in (**A**) lab and (**B**) wild mouse gut microbiota. *Esch/Shig* = Escherichia/Shigella.

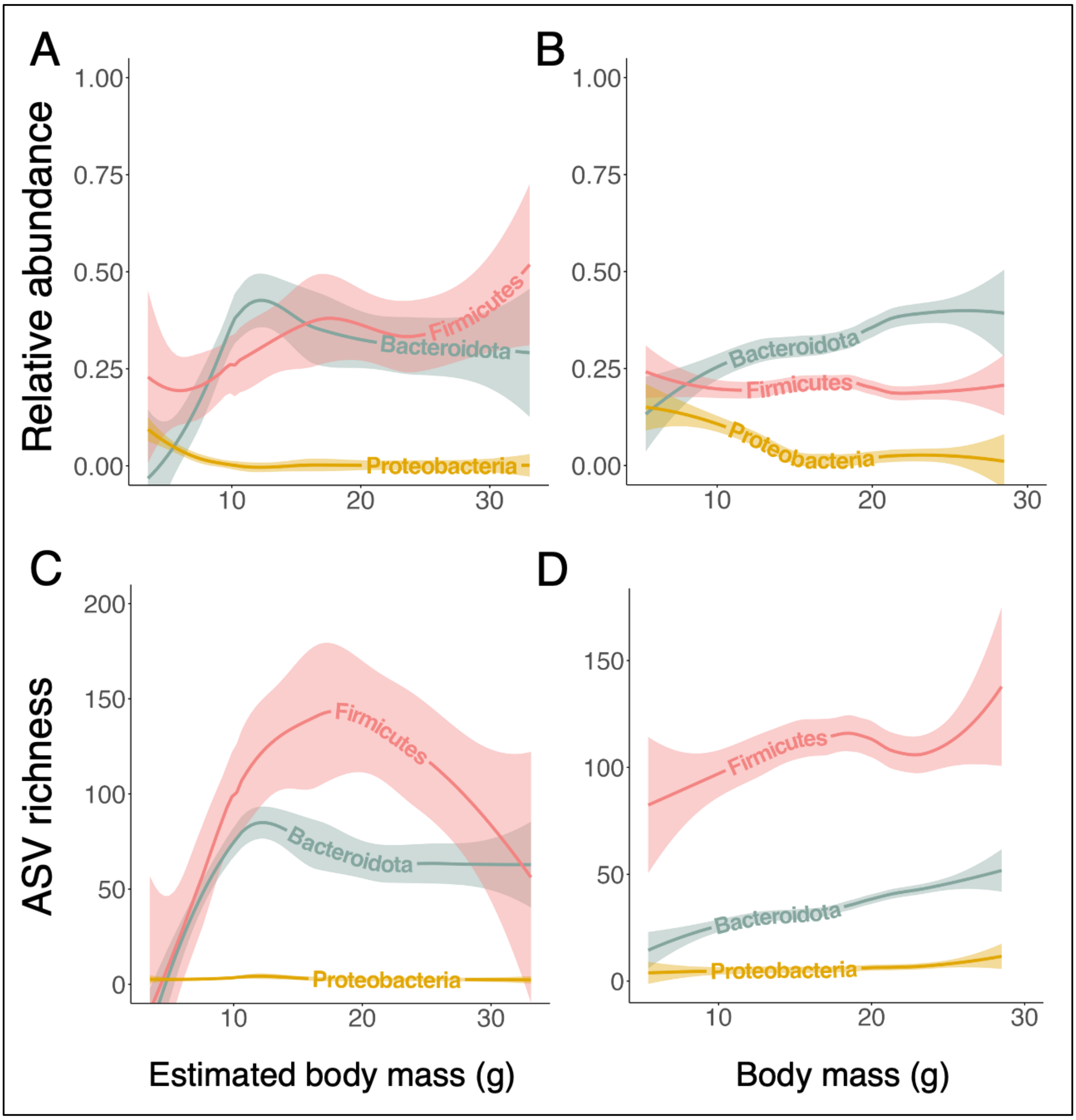


**Supplementary Figure 6.** Dynamics in bacterial phyla based on amplicon sequence variants (ASVs) that were only detected in (**A**, **C**) lab or (**B**, **D**) wild mice. (**A–B**) Relative abundance of Firmicutes, Bacteroidota, and Proteobacteria. (**C–D**) ASV richness (total count of unique ASVs) in Firmicutes, Bacteroidota, and Proteobacteria. Lines are locally estimated scatterplot smoothing (LOESS) lines with 95% confidence interval bands.

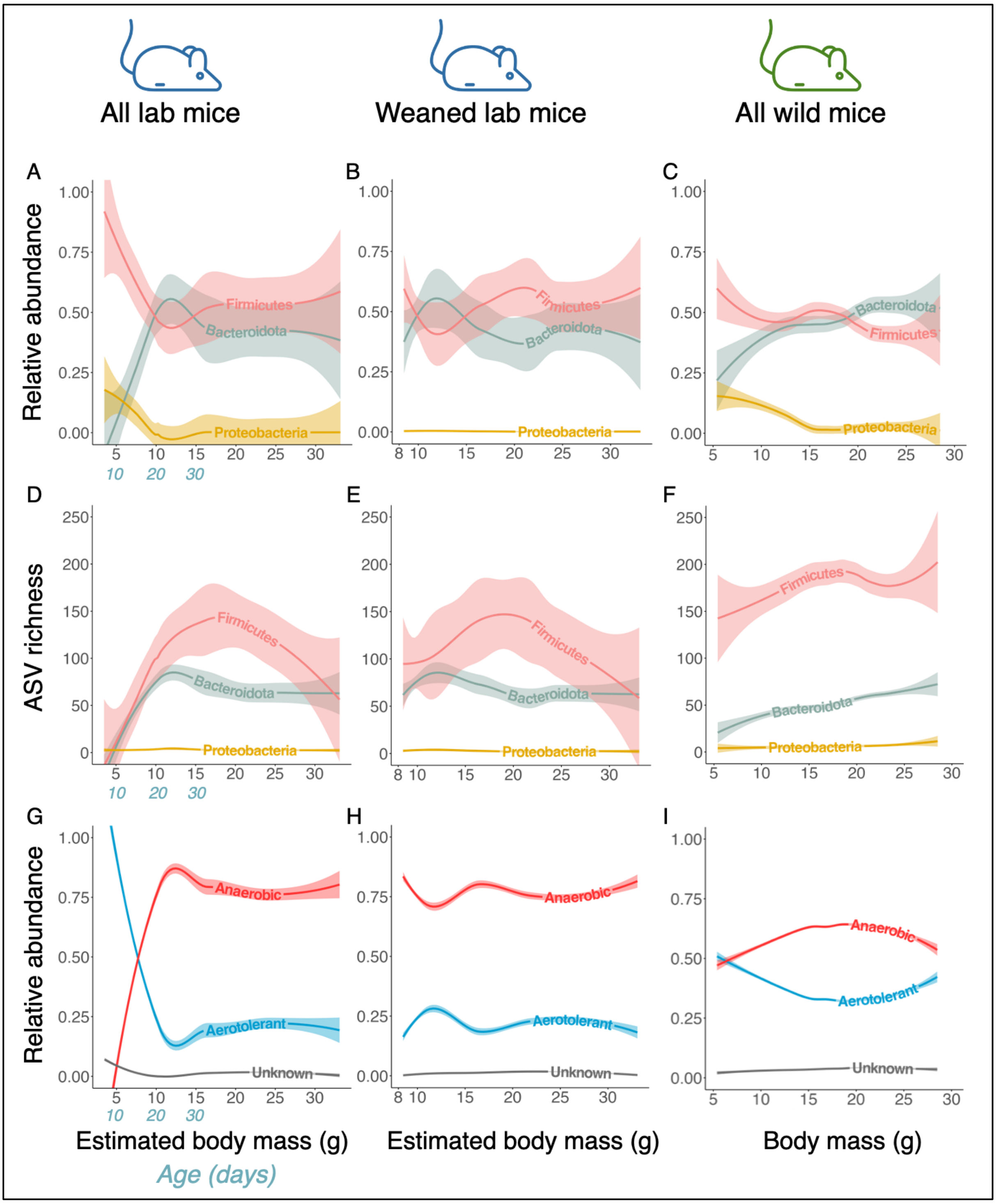


**Supplementary Figure 7.** Age-related gut microbial dynamics in (**A, B, D, E, G, H**) lab and (**C, F, I**) wild mice. Lab mouse data is limited to samples from weaned individuals in middle panels (**B, E, H**). Relative abundance of (**A–C**) predominant phyla (abundances were measured from all taxa), (**D–F**) ASV richness (total count of unique ASVs) in predominant phyla, and (**G–I**) aerotolerant and obligate anaerobic bacteria across (estimated) body mass. Lines are locally estimated scatterplot smoothing (LOESS) lines with 95% confidence interval bands. *Anaerobic* = obligate anaerobes, *aerotolerant* = everything else with known aerotolerance, *unknown* = bacteria with unknown aerotolerance.

**Supplementary Table 1. Aerotolerance and spore-forming ability of bacterial genera detected across laboratory and wild mice.** Bergey's Manual of Systematics of Archae and Bacteria alongside additional references (listed below) were used to determine aerotolerance (A = aerotolerant, OA = obligate anaerobe). Bacteria were classified as obligate anaerobes **only** when explicitly listed as *obligate* anaerobes. Aerotolerance was determined based on genus listed in *Genus*. If multiple genera were assigned for a given ASV (e.g., "Methylobacterium-Methylorubrum", genus based on which aerotolerance was determined is listed in *Comments*. If genus-level information was not available, family-level information was inspected and used if all genera from a given family were stated to have same aerotolerance (indicated in *Comments* as 'Based on -ceae').

*Supplementary Table is provided separately*

### Supplementary references


1. Spangenberg, E., Wallenbeck, A., Eklöf, A.-C., Carlstedt-Duke, J. & Tjäder, S. Housing breeding mice in three different IVC systems: maternal performance and pup development. *Lab Anim* **48**, 193–206 (2014).
2. Body Weight Information for C57BL/6J | The Jackson Laboratory. https://www.jax.org/jax-mice-and-services/strain-data-sheet-pages/body-weight-chart-000664.
3. Gray, M. M. *et al.* Genetics of Rapid and Extreme Size Evolution in Island Mice. *Genetics* **201**, 213–228 (2015).
4. Ferrari, M., Lindholm, A. K. & König, B. The risk of exploitation during communal nursing in house mice, Mus musculus domesticus. *Anim Behav* **110**, 133–143 (2015).
