## Supplementary Table 1 for "Early-life gut microbiota assembly patterns are conserved between laboratory and wild mice"

| Aerotolerance_category | |
| --- | --- |
| A | Aerotolerant |
| AN | Anaerobic |
| N/A | Not applicable |

| Family | Genus | Aerotolerance_category | Reference | Comments |
| --- | --- | --- | --- | --- |
| [Clostridium] methylpentosum group | Other | N/A | NA | |
| [Clostridium] methylpentosum group | Other | N/A | Bergey's manual | Varies within family |
| [Eubacterium] coprostanoligenes group | Other | AN | Bergey's manual | Based on Eubacterium |
| [Eubacterium] coprostanoligenes group | Other | N/A | NA | |
| 67-14 | Other | N/A | NA | |
| A4b | Other | N/A | NA | |
| Abditibacteriaceae | Abditibacterium | N/A | NA | |
| Acetobacteraceae | Acetobacter | A | Bergey's Manual | |
| Acetobacteraceae | Acidicaldus | A | Johnson et al., 2006 | |
| Acetobacteraceae | Acidiphilium | A | Bergey's Manual | |
| Acetobacteraceae | Acidisoma | A | Bergey's Manual | |
| Acetobacteraceae | Endobacter | A | Bergey's Manual | Based on Acetobacteraceae |
| Acetobacteraceae | Gluconobacter | A | Bergey's Manual | |
| Acetobacteraceae | Other | A | Bergey's Manual | Based on Acetobacteraceae |
| Acetobacteraceae | Other | A | Bergey's manual | |
| Acetobacteraceae | Rhodovastum | A | Bergey's Manual | Based on Acetobacteraceae |
| Acetobacteraceae | Roseomonas | A | Bergey's Manual | |
| Acholeplasmataceae | Anaeroplasma | AN | Bergey's Manual | |
| Acholeplasmataceae | Other | N/A | NA | |
| Acholeplasmataceae | Other | N/A | | |
| Acidaminococcaceae | Phascolarctobacterium | AN | Bergey's Manual | |
| Acidobacteriaceae (Subgroup 1) | Edaphobacter | A | Bergey's Manual | |
| Acidothermaceae | Acidothermus | A | Bergey's Manual | |
| Actinomycetaceae | Actinomyces | A | Bergey's Manual | |
| Aerococcaceae | Facklamia | A | Bergey's Manual | |
| Aerococcaceae | Other | A | Bergey's Manual | Based on Aerococcaceae |
| Aerococcaceae | Other | A | Bergey's manual | |
| Aeromonadaceae | Oceanisphaera | A | Bergey's Manual | Based on Aeromonadaceae |
| Akkermansiaceae | Akkermansia | AN | Bergey's Manual | |
| AKYG1722 | Other | N/A | NA | |
| Alcaligenaceae | Achromobacter | A | Bergey's Manual | |
| Alcaligenaceae | Alcaligenes | A | Bergey's Manual | Based on Alcaligenaceae |
| Alcaligenaceae | Candidimonas | A | Bergey's Manual | |
| Alcaligenaceae | Eoetvoesia | A | Bergey's Manual | Based on Alcaligenaceae |
| Alcaligenaceae | Other | A | Bergey's Manual | Based on Alcaligenaceae |

| | | | | |
|---|---|---|---|---|
| Alcaligenaceae | Other | A | Bergey's manual | |
| Alcaligenaceae | Paenalcaligenes | A | Bergey's Manual | |
| Alcaligenaceae | Verticiella | A | Vandamme et al. 2015 | Based on Verticia |
| Alicyclobacillaceae | Tumebacillus | A | Bergey's Manual | |
| Amoebophilaceae | Candidatus Cardinium | N/A | | |
| Anaerofustaceae | Anaerofustis | AN | Bergey's Manual | |
| Anaerovoracaceae | [Eubacterium] brachy gro | AN | Bergey's Manual | Based on Eubacterium |
| Anaerovoracaceae | [Eubacterium] nodatum g | AN | Bergey's Manual | Based on Eubacterium |
| Anaerovoracaceae | Anaerovorax | AN | Bergey's Manual | |
| Anaerovoracaceae | Family XIII AD3011 group | A | Bergey's Manual | Based on Anaerovorax |
| Anaerovoracaceae | Family XIII UCG-001 | A | Bergey's Manual | Based on Anaerovorax |
| Anaerovoracaceae | Other | N/A | NA | |
| Anaerovoracaceae | Other | N/A | | |
| Anaplasmataceae | Wolbachia | N/A | | |
| Atopobiaceae | Coriobacteriaceae UCG-0 | A | Bergey's Manual | Based on Atopobiaceae |
| Atopobiaceae | Coriobacteriaceae UCG-0 | A | Bergey's Manual | Based on Atopobiaceae |
| Azospirillaceae | Skermanella | A | Bergey's Manual | |
| Bacillaceae | Allobacillus | N/A | | |
| Bacillaceae | Bacillus | A | Bergey's Manual | |
| Bacillaceae | Falsibacillus | N/A | | |
| Bacillaceae | Microaerobacter | N/A | | |
| Bacillaceae | Natronobacillus | N/A | | |
| Bacillaceae | Other | A | Bergey's manual | |
| Bacillaceae | Other | N/A | NA | |
| Bacteroidaceae | Bacteroides | A | Bergey's Manual | |
| Bacteroidales RF16 group | Other | A | Bergey's Manual | Based on Bacteroidales RF16 group |
| Bacteroidales RF16 group | Other | N/A | | |
| Barnesiellaceae | Barnesiella | AN | Bergey's Manual | |
| Barnesiellaceae | Coprobacter | N/A | | |
| Barnesiellaceae | Other | N/A | NA | |
| Barnesiellaceae | Other | N/A | | |
| Bdellovibrionaceae | Bdellovibrio | A | Bergey's Manual | |
| Beijerinckiaceae | 1174-901-12 | N/A | NA | |
| Beijerinckiaceae | Bosea | A | Bergey's Manual | |
| Beijerinckiaceae | Methylobacterium-Methy | A | Bergey's Manual | Based on Methylobacterium |
| Beijerinckiaceae | Microvirga | A | Kanso & Patel 2003 | |
| Beijerinckiaceae | Other | A | Bergey's Manual | Based on Beijerinckiaceae |
| Beijerinckiaceae | Other | A | Bergey's manual | |
| Beijerinckiaceae | Roseiarcus | N/A | | |
| Bifidobacteriaceae | Bifidobacterium | A | Bergey's Manual | |
| Bradymonadaceae | Bradymonas | N/A | | |

| | | | | |
|---|---|---|---|---|
| Brevibacteriaceae | Brevibacterium | A | Bergey's Manual | |
| Brevibacteriaceae | Spelaeicoccus | N/A | | |
| Bryobacteraceae | Bryobacter | A | Bergey's Manual | |
| Budviciaceae | Budvicia | A | Bergey's Manual | |
| Butyricicoccaceae | Butyricicoccus | AN | Trachsel et al., 2018 | |
| Butyricicoccaceae | Other | N/A | NA | |
| Butyricicoccaceae | Other | N/A | | |
| Butyricicoccaceae | UCG-008 | N/A | NA | |
| Butyricicoccaceae | UCG-009 | AN | Bergey's Manual | Based on Clostridia |
| Caldicoprobacteraceae | Caldicoprobacter | AN | Bergey's Manual | |
| Candidatus Hepatincola | Other | N/A | NA | |
| Candidatus Hepatincola | Other | N/A | | |
| Carnobacteriaceae | Atopostipes | A | Bergey's Manual | |
| Carnobacteriaceae | Carnobacterium | A | Bergey's Manual | |
| Carnobacteriaceae | Granulicatella | A | Bergey's Manual | |
| Carnobacteriaceae | Marinilactibacillus | A | Bergey's Manual | |
| Catellicoccaceae | Catellicoccus | N/A | | |
| Caulobacteraceae | Brevundimonas | A | Bergey's Manual | |
| Caulobacteraceae | Caulobacter | A | Bergey's Manual | |
| Caulobacteraceae | Phenylobacterium | A | Bergey's Manual | |
| Cellulomonadaceae | Cellulomonas | A | Bergey's Manual | |
| Cellulomonadaceae | Oerskovia | A | Bergey's Manual | |
| Cellulomonadaceae | Paraoerskovia | A | Bergey's Manual | |
| Cellulomonadaceae | Pseudactinotalea | N/A | | |
| Cellvibrionaceae | Cellvibrio | A | Bergey's Manual | |
| Chitinophagaceae | Chitinophaga | A | Bergey's Manual | Based on Chitinophagaceae |
| Chitinophagaceae | Haoranjiania | A | Bergey's Manual | |
| Chitinophagaceae | Other | A | Bergey's Manual | |
| Chitinophagaceae | Other | A | Bergey's manual | |
| Chitinophagaceae | Segetibacter | A | Bergey's Manual | |
| Christensenellaceae | Christensenella | AN | Morotomi et al., 2012 | |
| Christensenellaceae | Christensenellaceae R-7 g | AN | Morotomi et al., 2012 | |
| Christensenellaceae | Other | AN | Morotomi et al., 2012 | |
| Christensenellaceae | Other | N/A | | |
| Chroococcidiopsaceae | Aliterella | N/A | | |
| Chthoniobacteraceae | Candidatus Udaeobacter | A | Bergey's Manual | |
| Clostridiaceae | Candidatus Arthromitus | AN | Schnupf et al., 2015 | |
| Clostridiaceae | Clostridium sensu stricto | AN | Bergey's Manual | Based on Clostridiaceae |
| Clostridiaceae | Clostridium sensu stricto | AN | Bergey's Manual | Based on Clostridiaceae |
| Clostridiaceae | Clostridium sensu stricto | AN | Bergey's Manual | Based on Clostridiaceae |
| Clostridiaceae | Clostridium sensu stricto | AN | Bergey's Manual | Based on Clostridiaceae |

| | | | | |
|---|---|---|---|---|
| Clostridiaceae | Other | AN | Bergey's Manual | Based on Clostridiaceae |
| Clostridiaceae | Other | AN | Bergey's manual | |
| Comamonadaceae | Comamonas | A | Bergey's Manual | |
| Comamonadaceae | Lampropedia | A | Bergey's Manual | |
| Comamonadaceae | Limnohabitans | A | Bergey's Manual | |
| Comamonadaceae | Other | N/A | Bergey's Manual | Based on Comamonadaceae |
| Comamonadaceae | Other | N/A | | |
| Comamonadaceae | Polaromonas | A | Bergey's Manual | |
| Comamonadaceae | Simplicispira | N/A | Bergey's Manual | Based on Comamonadaceae |
| Comamonadaceae | Variovorax | A | Bergey's Manual | |
| Comamonadaceae | Xenophilus | N/A | Bergey's Manual | Based on Comamonadaceae |
| Coralloluteibacterium | Other | N/A | NA | |
| Coriobacteriaceae | Collinsella | AN | Bergey's Manual | |
| Coriobacteriales Incertae Sedis | Other | N/A | NA | |
| Corynebacteriaceae | Corynebacterium | A | Bergey's Manual | |
| Corynebacteriales Incertae Sedis | Tomitella | N/A | | |
| Crocinitomicaceae | Fluviicola | A | Bergey's Manual | |
| Crocinitomicaceae | Other | N/A | NA | |
| Crocinitomicaceae | Other | N/A | | |
| Cryomorphaceae | Other | A | Bergey's Manual | Based on Cryomorphaceae |
| Cryomorphaceae | Other | A | Bergey's manual | |
| Cyanobiaceae | Synechococcus CC9902 | N/A | | |
| Cyclobacteriaceae | Algoriphagus | A | Bergey's Manual | |
| Cyclobacteriaceae | Other | A | Bergey's Manual | Based on Cyclobacteriaceae |
| Cyclobacteriaceae | Other | A | Bergey's manual | |
| Deferribacteraceae | Mucispirillum | A | Bergey's Manual | Based on Deferribacteraceae |
| Defluviitaleaceae | Defluviitaleaceae UCG-01 | A | Bergey's Manual | Based on Defluviitaleaceae |
| Demequinaceae | Demequina | N/A | | |
| Dermabacteraceae | Brachybacterium | A | Bergey's Manual | |
| Dermabacteraceae | Helcobacillus | N/A | Bergey's Manual | Based on Dermabacteraceae |
| Dermacoccaceae | Flexivirga | A | Bergey's Manual | Based on Dermacoccaceae |
| Dermacoccaceae | Other | A | Bergey's Manual | Based on Dermacoccaceae |
| Desulfovibrionaceae | Bilophila | AN | Bergey's Manual | |
| Desulfovibrionaceae | Desulfovibrio | AN | Bergey's Manual | |
| Desulfovibrionaceae | Lawsonia | AN | Bergey's Manual | Based on Desulfovibrionaceae |
| Desulfovibrionaceae | Other | AN | Bergey's Manual | Based on Desulfovibrionaceae |
| DEV007 | Other | N/A | NA | |
| Devosiaceae | Arsenicitalea | N/A | | |
| Devosiaceae | Devosia | A | Bergey's Manual | |
| Devosiaceae | Other | N/A | NA | |
| Devosiaceae | Other | N/A | | |

| | | | | |
|---|---|---|---|---|
| Devosiaceae | Pelagibacterium | N/A | | |
| Dietziaceae | Dietzia | A | Bergey's Manual | |
| Diplorickettsiaceae | Diplorickettsia | A | Mediannikov et al., 2010 | |
| Diplorickettsiaceae | Other | N/A | NA | |
| Diplorickettsiaceae | Other | N/A | | |
| Diplorickettsiaceae | Rickettsiella | A | Bergey's Manual | |
| Dysgonomonadaceae | Dysgonomonas | A | Bergey's Manual | |
| Eggerthellaceae | Adlercreutzia | AN | Bergey's Manual | |
| Eggerthellaceae | Asaccharobacter | A | Bergey's Manual | Based on Eggerthellaceae |
| Eggerthellaceae | DNF00809 | N/A | Bergey's Manual | Based on Eggerthellaceae |
| Eggerthellaceae | Eggerthella | AN | Bergey's Manual | |
| Eggerthellaceae | Enterorhabdus | A | Clavel et al., 2009 | |
| Eggerthellaceae | Gordonibacter | AN | Wurdemann et al., 2009, Bergey's Manual | |
| Eggerthellaceae | Other | A | Bergey's Manual | Based on Eggerthellaceae |
| Eggerthellaceae | Parvibacter | A | Clavel et al., 2013 | |
| Enterobacteriaceae | Aquamonas | A | Bergey's Manual, Degelmann et a | Based on Enterobacteriaceae |
| Enterobacteriaceae | Atlantibacter | A | Bergey's Manual, Degelmann et a | Based on Enterobacteriaceae |
| Enterobacteriaceae | Buttiauxella | A | Bergey's Manual, Degelmann et al., 2009 | |
| Enterobacteriaceae | Cedecea | A | Bergey's Manual, Degelmann et al., 2010 | |
| Enterobacteriaceae | Citrobacter | A | Bergey's Manual, Degelmann et al., 2011 | |
| Enterobacteriaceae | Escherichia-Shigella | A | Bergey's Manual, Degelmann et a | Based on Escherichia |
| Enterobacteriaceae | Klebsiella | A | Bergey's Manual, Degelmann et al., 2013 | |
| Enterobacteriaceae | Kluyvera | A | Bergey's Manual, Degelmann et al., 2014 | |
| Enterobacteriaceae | Kosakonia | A | Bergey's Manual, Degelmann et a | Based on Enterobacteriaceae |
| Enterobacteriaceae | Other | A | Bergey's Manual, Degelmann et a | Based on Enterobacteriaceae |
| Enterobacteriaceae | Other | A | Bergey's Manual, Degelmann et al., 2009 | |
| Enterobacteriaceae | Raoultella | A | Bergey's Manual, Degelmann et a | Based on Enterobacteriaceae |
| Enterobacteriaceae | Salmonella | A | Bergey's Manual, Degelmann et al., 2015 | |
| Enterobacteriaceae | Yokenella | A | Bergey's Manual, Degelmann et al., 2016 | |
| Enterococcaceae | Enterococcus | A | Bergey's Manual | |
| Enterococcaceae | Other | N/A | NA | |
| Enterococcaceae | Other | N/A | Bergey's manual | N/A aerotolerance |
| Entomoplasmatales Incertae Sedis | Candidatus Hepatoplasm | N/A | | |
| Erwiniaceae | Erwinia | A | Bergey's Manual | |
| Erwiniaceae | Other | N/A | NA | |
| Erwiniaceae | Other | N/A | | |
| Erwiniaceae | Pantoea | A | Bergey's Manual | |
| Erwiniaceae | Siccibacter | N/A | | |
| Erwiniaceae | Tatumella | A | Bergey's Manual | |
| Erysipelatoclostridiaceae | Candidatus Stoquefichus | A | Bergey's Manual | Based on Erysipelatoclostridiaceae |
| Erysipelatoclostridiaceae | Catenibacterium | AN | Bergey's Manual | |

| | | | | |
|---|---|---|---|---|
| Erysipelatoclostridiaceae | Coprobacillus | AN | Bergey's Manual | |
| Erysipelatoclostridiaceae | Erysipelatoclostridium | AN | Yutin & Galperin, 2013 | |
| Erysipelatoclostridiaceae | Erysipelotrichaceae UCG- | A | Bergey's Manual | Based on Erysipelatoclostridiaceae |
| Erysipelatoclostridiaceae | Other | A | Bergey's Manual | Based on Erysipelatoclostridiaceae |
| Erysipelatoclostridiaceae | Other | N/A | | |
| Erysipelotrichaceae | [Clostridium] innocuum g | A | Bergey's Manual | Based on Erysipelotrichaceae |
| Erysipelotrichaceae | Dubosiella | AN | Cox et al., 2017 | |
| Erysipelotrichaceae | Erysipelotrichaceae UCG- | A | Bergey's Manual | Based on Erysipelotrichaceae |
| Erysipelotrichaceae | Faecalibaculum | AN | Chang et al., 2015 | |
| Erysipelotrichaceae | Faecalitalea | A | Bergey's Manual | Based on Erysipelotrichaceae |
| Erysipelotrichaceae | Holdemanella | A | Bergey's Manual | Based on Erysipelotrichaceae |
| Erysipelotrichaceae | Ileibacterium | A | Bergey's Manual | Based on Erysipelotrichaceae |
| Erysipelotrichaceae | Other | A | Bergey's Manual | Based on Erysipelotrichaceae |
| Erysipelotrichaceae | Other | A | Bergey's manual | |
| Erysipelotrichaceae | Turicibacter | A | Bergey's Manual | |
| Erysipelotrichaceae | ZOR0006 | A | Bergey's Manual | Based on Erysipelotrichaceae |
| Eubacteriaceae | Other | AN | Bergey's Manual | Based on Eubacteriaceae |
| Eubacteriaceae | Other | AN | Bergey's manual | |
| Euzebyaceae | Other | N/A | NA | |
| Exiguobacteraceae | Exiguobacterium | A | Bergey's Manual | |
| Family XI | Other | N/A | NA | |
| Family XI | Tissierella | AN | Bergey's Manual | |
| Flavobacteriaceae | Aequorivita | A | Bergey's Manual | |
| Flavobacteriaceae | Aquibacter | A | Bergey's Manual | Based on Flavobacteriaceae |
| Flavobacteriaceae | Arenibacter | A | Bergey's Manual | |
| Flavobacteriaceae | Aurantiacicella | A | Bergey's Manual | Based on Flavobacteriaceae |
| Flavobacteriaceae | Flavobacterium | A | Bergey's Manual | |
| Flavobacteriaceae | Gelidibacter | A | Bergey's Manual | |
| Flavobacteriaceae | Gillisia | A | Bergey's Manual | |
| Flavobacteriaceae | Imtechella | A | Bergey's Manual | Based on Flavobacteriaceae |
| Flavobacteriaceae | Leeuwenhoekiella | A | Bergey's Manual | |
| Flavobacteriaceae | Muricauda | A | Bergey's Manual | |
| Flavobacteriaceae | Myroides | A | Bergey's Manual | |
| Flavobacteriaceae | Other | A | Bergey's Manual | Based on Flavobacteriaceae |
| Flavobacteriaceae | Other | N/A | Bergey's manual | |
| Flavobacteriaceae | Subsaxibacter | A | Bergey's Manual | |
| Frankiaceae | Frankia | A | Bergey's Manual | |
| Frankiaceae | Jatrophihabitans | A | Bergey's Manual | |
| Fusobacteriaceae | Cetobacterium | A | Bergey's Manual | |
| Fusobacteriaceae | Fusobacterium | AN | Bergey's Manual | |
| Gaiellaceae | Gaiella | A | Bergey's Manual | |

| | | | | |
|---|---|---|---|---|
| Garciellaceae | Rhabdanaerobium | AN | Bergey's Manual | |
| Gemellaceae | Gemella | A | Bergey's Manual | |
| Geminicoccaceae | Candidatus Alysiosphaera | N/A | | |
| Geminicoccaceae | Geminicoccus | N/A | | |
| Geminicoccaceae | Other | N/A | NA | |
| Geminicoccaceae | Other | N/A | | |
| Gemmataceae | Fimbriiglobus | A | Bergey's Manual | |
| Gemmataceae | Gemmata | A | Bergey's Manual | |
| Gemmataceae | Other | A | Bergey's Manual | Based on Gemmataceae |
| Gemmatimonadaceae | Gemmatimonas | A | Bergey's Manual | |
| Gemmatimonadaceae | Other | N/A | NA | |
| Geodermatophilaceae | Antricoccus | A | Bergey's Manual | Based on Geodermatophilaceae |
| Geodermatophilaceae | Blastococcus | A | Bergey's Manual | |
| Geodermatophilaceae | Klenkia | A | Bergey's Manual | |
| Geodermatophilaceae | Modestobacter | A | Bergey's Manual | |
| Gottschalkia | Other | N/A | NA | |
| Gottschalkia | Other | N/A | | |
| Granulosicoccaceae | Granulosicoccus | N/A | | |
| Hafniaceae | Edwardsiella | A | Bergey's Manual | |
| Hafniaceae | Hafnia-Obesumbacterium | A | Bergey's Manual | Based on Hafniaceae |
| Halomonadaceae | Chromohalobacter | A | Bergey's Manual | |
| Halomonadaceae | Halomonas | A | Bergey's Manual | |
| Halomonadaceae | Salinicola | A | Bergey's Manual | |
| Helicobacteraceae | Helicobacter | A | Bergey's Manual | |
| Hydrogenoanaerobacterium | Other | AN | Bergey's manual | |
| Hydrogenoanaerobacterium | Other | N/A | NA | |
| Hyphomicrobiaceae | Hyphomicrobium | A | Bergey's Manual | |
| Hyphomicrobiaceae | Pedomicrobium | A | Bergey's Manual | |
| Iamiaceae | Iamia | A | Bergey's Manual | |
| Ilumatobacteraceae | CL500-29 marine group | N/A | | |
| Ilumatobacteraceae | Ilumatobacter | N/A | | |
| Ilumatobacteraceae | Other | N/A | NA | |
| Intrasporangiaceae | Humibacillus | A | Bergey's Manual | |
| Intrasporangiaceae | Intrasporangium | A | Bergey's Manual | |
| Intrasporangiaceae | Janibacter | A | Bergey's Manual | |
| Intrasporangiaceae | Knoellia | A | Bergey's Manual | |
| Intrasporangiaceae | Oryzihumus | A | Bergey's Manual | |
| Intrasporangiaceae | Other | A | Bergey's Manual | Based on Intrasporangiaceae |
| Intrasporangiaceae | Pedococcus-Phycicoccus | A | Bergey's Manual | Based on Intrasporangiaceae |
| Isosphaeraceae | Aquisphaera | A | Bergey's Manual | |
| Isosphaeraceae | Candidatus Nostocoida | A | Bergey's Manual | Based on Isosphaeraceae |

| | | | | |
|---|---|---|---|---|
| Isosphaeraceae | Isosphaera | A | Bergey's Manual | |
| Isosphaeraceae | Other | A | Bergey's Manual | Based on Isosphaeraceae |
| Isosphaeraceae | Paludisphaera | A | Bergey's Manual | |
| Isosphaeraceae | Singulisphaera | A | Bergey's Manual | |
| Isosphaeraceae | Tundrisphaera | A | Bergey's Manual | Based on Isosphaeraceae |
| JG30-KF-CM45 | Other | N/A | NA | |
| Kaistiaceae | Kaistia | N/A | | |
| Kineosporiaceae | Angustibacter | A | Bergey's Manual | |
| Kineosporiaceae | Other | N/A | NA | |
| Kineosporiaceae | Quadrisphaera | N/A | Bergey's Manual | |
| Ktedonobacteraceae | G12-WMSP1 | A | Bergey's Manual | Based on Ktedonobacteraceae |
| Ktedonobacteraceae | HSB OF53-F07 | A | Bergey's Manual | Based on Ktedonobacteraceae |
| Labraceae | Labrys | A | Bergey's Manual | |
| Lachnospiraceae | 28-Apr | AN | Bergey's Manual | Based on Lachnospiraceae |
| Lachnospiraceae | [Acetivibrio] ethanolgigne | AN | Bergey's Manual | Based on Lachnospiraceae |
| Lachnospiraceae | [Eubacterium] eligens gro | AN | Bergey's Manual | Based on Lachnospiraceae |
| Lachnospiraceae | [Eubacterium] fissicatena | AN | Bergey's Manual | Based on Lachnospiraceae |
| Lachnospiraceae | [Eubacterium] hallii grou | AN | Bergey's Manual | Based on Lachnospiraceae |
| Lachnospiraceae | [Eubacterium] oxidoreduc | AN | Bergey's Manual | Based on Lachnospiraceae |
| Lachnospiraceae | [Eubacterium] ventriosun | AN | Bergey's Manual | Based on Lachnospiraceae |
| Lachnospiraceae | [Eubacterium] xylanophil | AN | Bergey's Manual | Based on Lachnospiraceae |
| Lachnospiraceae | [Ruminococcus] gauvreau | AN | Bergey's Manual | Based on Lachnospiraceae |
| Lachnospiraceae | [Ruminococcus] gnavus g | AN | Bergey's Manual | Based on Lachnospiraceae |
| Lachnospiraceae | [Ruminococcus] torques g | AN | Bergey's Manual | Based on Lachnospiraceae |
| Lachnospiraceae | A2 | AN | Bergey's Manual | Based on Lachnospiraceae |
| Lachnospiraceae | Acetatifactor | AN | Bergey's Manual | Based on Lachnospiraceae |
| Lachnospiraceae | Agathobacter | AN | Bergey's Manual | Based on Lachnospiraceae |
| Lachnospiraceae | Anaerosporobacter | AN | Bergey's Manual | Based on Lachnospiraceae |
| Lachnospiraceae | Anaerostipes | AN | Bergey's Manual | Based on Lachnospiraceae |
| Lachnospiraceae | ASF356 | AN | Bergey's Manual | Based on Lachnospiraceae |
| Lachnospiraceae | Blautia | AN | Bergey's Manual | Based on Lachnospiraceae |
| Lachnospiraceae | Cellulosilyticum | AN | Bergey's Manual | Based on Lachnospiraceae |
| Lachnospiraceae | Coprococcus | AN | Bergey's Manual | Based on Lachnospiraceae |
| Lachnospiraceae | Cuneatibacter | AN | Bergey's Manual | Based on Lachnospiraceae |
| Lachnospiraceae | Dorea | AN | Bergey's Manual | Based on Lachnospiraceae |
| Lachnospiraceae | Eisenbergiella | AN | Bergey's Manual | Based on Lachnospiraceae |
| Lachnospiraceae | Epulopiscium | AN | Bergey's Manual | Based on Lachnospiraceae |
| Lachnospiraceae | Fusicatenibacter | AN | Bergey's Manual | Based on Lachnospiraceae |
| Lachnospiraceae | GCA-900066575 | AN | Bergey's Manual | Based on Lachnospiraceae |
| Lachnospiraceae | Hungatella | AN | Bergey's Manual | Based on Lachnospiraceae |
| Lachnospiraceae | Lachnoclostridium | AN | Bergey's Manual | Based on Lachnospiraceae |

| | | | | |
|---|---|---|---|---|
| Lachnospiraceae | Lachnospiraceae FCS020 | AN | Bergey's Manual | Based on Lachnospiraceae |
| Lachnospiraceae | Lachnospiraceae NK4A13 | AN | Bergey's Manual | Based on Lachnospiraceae |
| Lachnospiraceae | Lachnospiraceae NK4B4 g | AN | Bergey's Manual | Based on Lachnospiraceae |
| Lachnospiraceae | Lachnospiraceae UCG-00 | AN | Bergey's Manual | Based on Lachnospiraceae |
| Lachnospiraceae | Lachnospiraceae UCG-00 | AN | Bergey's Manual | Based on Lachnospiraceae |
| Lachnospiraceae | Lachnospiraceae UCG-00 | AN | Bergey's Manual | Based on Lachnospiraceae |
| Lachnospiraceae | Lachnospiraceae UCG-00 | AN | Bergey's Manual | Based on Lachnospiraceae |
| Lachnospiraceae | Lachnospiraceae UCG-00 | AN | Bergey's Manual | Based on Lachnospiraceae |
| Lachnospiraceae | Lachnospiraceae UCG-01 | AN | Bergey's Manual | Based on Lachnospiraceae |
| Lachnospiraceae | Marvinbryantia | AN | Bergey's Manual | Based on Lachnospiraceae |
| Lachnospiraceae | Murimonas | AN | Bergey's Manual | Based on Lachnospiraceae |
| Lachnospiraceae | Other | AN | Bergey's Manual | Based on Lachnospiraceae |
| Lachnospiraceae | Other | AN | Bergey's manual | |
| Lachnospiraceae | possible genus Sk018 | AN | Bergey's Manual | Based on Lachnospiraceae |
| Lachnospiraceae | Robinsoniella | AN | Bergey's Manual | Based on Lachnospiraceae |
| Lachnospiraceae | Roseburia | AN | Bergey's Manual | Based on Lachnospiraceae |
| Lachnospiraceae | Sellimonas | AN | Bergey's Manual | Based on Lachnospiraceae |
| Lachnospiraceae | Tuzzerella | AN | Bergey's Manual | Based on Lachnospiraceae |
| Lachnospiraceae | Tyzzerella | AN | Bergey's Manual | Based on Lachnospiraceae |
| Lactobacillaceae | Agrilactobacillus | N/A | Bergey's Manual | Based on Lactobacillaceae |
| Lactobacillaceae | Bombilactobacillus | N/A | Bergey's Manual | Based on Lactobacillaceae |
| Lactobacillaceae | Companilactobacillus | N/A | Bergey's Manual | Based on Lactobacillaceae |
| Lactobacillaceae | Dellaglioa | N/A | Bergey's Manual | Based on Lactobacillaceae |
| Lactobacillaceae | HT002 | N/A | Bergey's Manual | Based on Lactobacillaceae |
| Lactobacillaceae | Lacticaseibacillus | N/A | Bergey's Manual | Based on Lactobacillaceae |
| Lactobacillaceae | Lactiplantibacillus | N/A | Bergey's Manual | Based on Lactobacillaceae |
| Lactobacillaceae | Lactobacillus | A | Bergey's Manual | |
| Lactobacillaceae | Latilactobacillus | N/A | Bergey's Manual | Based on Lactobacillaceae |
| Lactobacillaceae | Leuconostoc | N/A | Bergey's Manual | |
| Lactobacillaceae | Levilactobacillus | N/A | Bergey's Manual | Based on Lactobacillaceae |
| Lactobacillaceae | Ligilactobacillus | A | Aerotolerance N/A based on Marta et al., 2021; Zheng et al., 2020 - so | |
| Lactobacillaceae | Limosilactobacillus | A | aerotolerance was N/A (Zheng et a., 2020) so BLASTed and blasts agai | |
| Lactobacillaceae | Other | N/A | Bergey's Manual | Based on Lactobacillaceae |
| Lactobacillaceae | Other | N/A | | |
| Lactobacillaceae | Paucilactobacillus | N/A | Bergey's Manual | Based on Lactobacillaceae |
| Lactobacillaceae | Weissella | A | Bergey's Manual | |
| Legionellaceae | Legionella | A | Bergey's Manual | |
| Leptolyngbyaceae | Leptolyngbya PCC-6306 | N/A | | |
| Listeriaceae | Listeria | A | Bergey's Manual | Based on Listeriaceae |
| Marinifilaceae | Butyricimonas | N/A | NA | |
| Marinifilaceae | Odoribacter | AN | Hardham et al., 2008 | |

| | | | | |
|---|---|---|---|---|
| Marinifilaceae | Other | N/A | NA | |
| Marinifilaceae | Other | N/A | | |
| Marinifilaceae | Sanguibacteroides | N/A | | |
| Marinilabiliaceae | Natronoflexus | N/A | Bergey's Manual | Based on Marinilabiliaceae |
| Marinilabiliaceae | Other | N/A | Bergey's Manual | Based on Marinilabiliaceae |
| Marinilabiliaceae | Other | N/A | | |
| Marinobacteraceae | Marinobacter | A | Bergey's Manual | |
| Methyloligellaceae | Methyloligella | N/A | | |
| Methyloligellaceae | Other | N/A | NA | |
| Methyloligellaceae | Other | N/A | | |
| Microbacteriaceae | Agrococcus | A | Bergey's Manual | |
| Microbacteriaceae | Amnibacterium | A | Bergey's Manual | |
| Microbacteriaceae | Curtobacterium | A | Bergey's Manual | |
| Microbacteriaceae | Frigoribacterium | A | Bergey's Manual | |
| Microbacteriaceae | Homoserinibacter | A | Bergey's Manual | |
| Microbacteriaceae | Leucobacter | N/A | Bergey's Manual | |
| Microbacteriaceae | Lysinimonas | A | Bergey's Manual | |
| Microbacteriaceae | Microbacterium | A | Bergey's Manual | |
| Microbacteriaceae | Microterricola | A | Bergey's Manual | |
| Microbacteriaceae | Mycetocola | A | Bergey's Manual | |
| Microbacteriaceae | Other | A | Bergey's Manual | Based on Microbacteriaceae |
| Microbacteriaceae | Parafrigoribacterium | A | Bergey's Manual | Based on Microbacteriaceae |
| Microbacteriaceae | Plantibacter | A | Bergey's Manual | |
| Microbacteriaceae | Pseudoclavibacter | A | Bergey's Manual | |
| Micrococcaceae | Arthrobacter | A | Bergey's Manual | |
| Micrococcaceae | Glutamicibacter | A | Bergey's Manual | |
| Micrococcaceae | Kocuria | A | Bergey's Manual | |
| Micrococcaceae | Nesterenkonia | A | Bergey's Manual | |
| Micrococcaceae | Other | N/A | NA | |
| Micrococcaceae | Paeniglutamicibacter | A | Bergey's Manual | |
| Micrococcaceae | Pseudarthrobacter | A | Bergey's Manual | |
| Micromonosporaceae | Actinoplanes | A | Bergey's Manual | |
| Micromonosporaceae | Micromonospora | A | Bergey's Manual | |
| Micromonosporaceae | Other | A | Bergey's Manual | Based on Micromonosporaceae |
| Micromonosporaceae | Xiangella | A | Bergey's Manual | |
| Monoglobaceae | Monoglobus | AN | Kim et al., 2017 | |
| Moraxellaceae | Acinetobacter | A | Bergey's Manual | |
| Moraxellaceae | Alkanindiges | A | Bergey's Manual | Based on Moraxellaceae |
| Moraxellaceae | Enhydrobacter | A | Bergey's Manual | |
| Moraxellaceae | Psychrobacter | A | Bergey's Manual | |
| Morganellaceae | Cosenzaea | N/A | | |

| | | | | |
|---|---|---|---|---|
| Morganellaceae | Moellerella | A | Bergey's Manual | |
| Morganellaceae | Morganella | A | Bergey's Manual | |
| Morganellaceae | Other | N/A | NA | |
| Morganellaceae | Other | N/A | | |
| Morganellaceae | Proteus | A | Bergey's Manual | |
| Morganellaceae | Providencia | A | Bergey's Manual | |
| Muribaculaceae | Muribaculum | AN | Bergey's Manual | |
| Muribaculaceae | Other | AN | Lagkouvardos et al 2019 | |
| Muribaculaceae | Other | AN | Lagkouvardos et al., 2019 | |
| MWH-CFBk5 | Other | N/A | NA | |
| MWH-CFBk5 | Other | N/A | | |
| Mycobacteriaceae | Mycobacterium | A | Bergey's Manual | |
| Mycobacteriaceae | Other | N/A | | |
| Mycoplasmataceae | Candidatus Bacilloplasma | A | Bergey's Manual | Based on Mycoplasmataceae |
| Mycoplasmataceae | Mycoplasma | N/A | Bergey's Manual | |
| Mycoplasmataceae | Other | A | Bergey's Manual | Based on Mycoplasmataceae |
| Mycoplasmataceae | Other | A | Bergey's manual | |
| Myxococcaceae | P3OB-42 | N/A | | |
| Nakamurellaceae | Nakamurella | A | Bergey's Manual | |
| Nannocystaceae | Enhygromyxa | N/A | | |
| Neisseriaceae | Other | A | Bergey's Manual | Based on Neisseriaceae |
| Neisseriaceae | Other | A | Bergey's manual | |
| Neisseriaceae | Vitreoscilla | A | Bergey's Manual | |
| Nitrospiraceae | Nitrospira | A | Bergey's Manual | |
| Nocardiaceae | Gordonia | A | Bergey's Manual | |
| Nocardiaceae | Nocardia | A | Bergey's Manual | |
| Nocardiaceae | Other | A | Bergey's Manual | Based on Nocardiaceae |
| Nocardiaceae | Rhodococcus | A | Bergey's Manual | |
| Nocardiaceae | Williamsia | A | Bergey's Manual | |
| Nocardioidaceae | Aeromicrobium | A | Bergey's Manual | |
| Nocardioidaceae | Marmoricola | A | Bergey's Manual | |
| Nocardioidaceae | Mumia | A | Bergey's Manual | Based on Nocardioidaceae |
| Nocardioidaceae | Nocardioides | A | Bergey's Manual | |
| Nocardioidaceae | Other | A | Bergey's Manual | Based on Nocardioidaceae |
| Nocardiopsaceae | Nocardiopsis | A | Bergey's Manual | |
| Nostocaceae | Calothrix PCC-6303 | N/A | | |
| Nostocaceae | Other | N/A | NA | |
| Nostocaceae | Rivularia PCC-7116 | N/A | Bergey's Manual | Based on Rivularia |
| Oligoflexaceae | Oligoflexus | N/A | | |
| Oscillospiraceae | Colidextribacter | AN | Ricaboni et al., 2017 | |
| Oscillospiraceae | Flavonifractor | AN | Bergey's Manual | Based on Eubacterium |

| | | | | |
|---|---|---|---|---|
| Oscillospiraceae | Intestinimonas | AN | Kläring et al., 2013, Bergey's manual | |
| Oscillospiraceae | NK4A214 group | AN | Tindal et al., 2019 | |
| Oscillospiraceae | Oscillibacter | AN | Iino et al. 2007 | |
| Oscillospiraceae | Oscillospira | AN | Bergey's Manual | |
| Oscillospiraceae | Other | A | Bergey's Manual | Based on Oscillospiraceae |
| Oscillospiraceae | Other | N/A | | |
| Oscillospiraceae | Papillibacter | AN | Bergey's Manual | |
| Oscillospiraceae | Pseudoflavonifractor | N/A | | |
| Oscillospiraceae | UCG-002 | N/A | NA | |
| Oscillospiraceae | UCG-003 | N/A | NA | |
| Oscillospiraceae | UCG-005 | AN | Bergey's Manual | Based on Oscillospiraceae |
| Oscillospiraceae | UCG-007 | N/A | NA | |
| Other | Other | N/A | NA | |
| Oxalobacteraceae | Duganella | A | Bergey's Manual | |
| Oxalobacteraceae | Massilia | A | Bergey's Manual | |
| Oxalobacteraceae | Noviherbaspirillum | A | Bergey's Manual | Based on Oxalobacteraceae |
| Oxalobacteraceae | Oxalicibacterium | A | Bergey's Manual | Based on Oxalobacteraceae |
| Oxalobacteraceae | Oxalobacter | AN | Bergey's Manual | |
| Oxalobacteraceae | Paraherbaspirillum | A | Bergey's Manual | Based on Oxalobacteraceae |
| Paenibacillaceae | Ammoniphilus | A | Bergey's Manual | |
| Paenibacillaceae | Paenibacillus | A | Bergey's Manual | |
| Paludibacteraceae | H1 | N/A | | |
| Pasteurellaceae | Conservatibacter | N/A | Bergey's Manual | |
| Pasteurellaceae | Mesocricetibacter | A | Bergey's Manual | |
| Pasteurellaceae | Muribacter | N/A | Bergey's Manual | |
| Pasteurellaceae | Rodentibacter | A | Bergey's Manual, Benga, Sager & Christensen 2018 | |
| Pectobacteriaceae | Nissabacter | N/A | | |
| Peptococcaceae | Other | AN | Bergey's Manual | Based on Peptococcaceae |
| Peptococcaceae | Other | AN | Bergey's manual | |
| Peptococcaceae | Peptococcus | AN | Bergey's Manual | |
| Peptostreptococcaceae | [Eubacterium] tenue grou | N/A | NA | |
| Peptostreptococcaceae | Clostridioides | A | Bergey's Manual | Based on Peptostreptococcaceae |
| Peptostreptococcaceae | Intestinibacter | A | Bergey's Manual | Based on Peptostreptococcaceae |
| Peptostreptococcaceae | Paeniclostridium | A | Bergey's Manual | Based on Peptostreptococcaceae |
| Peptostreptococcaceae | Paraclostridium | A | Bergey's Manual | |
| Peptostreptococcaceae | Romboutsia | AN | Gerritsen et al., BioRxiv 2019 | |
| Peptostreptococcaceae | Sporacetigenium | AN | Bergey's Manual | |
| Peptostreptococcaceae | Terrisporobacter | A | Bergey's Manual | Based on Peptostreptococcaceae |
| Phormidesmiaceae | Phormidesmis ANT.LACV | N/A | | |
| Phormidiaceae | Other | N/A | NA | |
| Phormidiaceae | Tychonema CCAP 1459-1 | N/A | NA | |

| | | | | |
|---|---|---|---|---|
| Pirellulaceae | Blastopirellula | A | Bergey's Manual | |
| Pirellulaceae | Bythopirellula | N/A | NA | |
| Pirellulaceae | Other | N/A | NA | |
| Pirellulaceae | Pir4 lineage | N/A | Bergey's Manual | Based on Pirellulaceae |
| Pirellulaceae | Pirellula | A | Bergey's Manual | |
| Pirellulaceae | Rhodopirellula | A | Bergey's Manual | |
| Pirellulaceae | Rubripirellula | A | Bergey's Manual | |
| Planococcaceae | Kurthia | A | Bergey's Manual | |
| Planococcaceae | Lysinibacillus | A | Bergey's Manual | |
| Planococcaceae | Other | A | Bergey's manual | |
| Planococcaceae | Other | N/A | | |
| Planococcaceae | Paenisporosarcina | A | Krishnamurthi et al., 2009 | |
| Planococcaceae | Planomicrobium | A | Bergey's Manual | |
| Planococcaceae | Psychrobacillus | A | Bergey's Manual | Based on Planococcaceae |
| Planococcaceae | Solibacillus | A | Krishnamurthi et al., 2009 | |
| Planococcaceae | Sporosarcina | A | Bergey's Manual | |
| Polyangiaceae | Aetherobacter | A | Garcia et al. | |
| Porphyromonadaceae | Falsiporphyromonas | AN | Bergey's Manual | Based on Porphyromonadaceae |
| Porphyromonadaceae | Other | AN | Bergey's Manual | Based on Porphyromonadaceae |
| Porphyromonadaceae | Other | AN | Bergey's manual | |
| Prevotellaceae | Alloprevotella | AN | Bergey's Manual | Based on Prevotellaceae |
| Prevotellaceae | Other | A | Bergey's Manual | Based on Prevotellaceae |
| Prevotellaceae | Other | AN | Bergey's manual | |
| Prevotellaceae | Paraprevotella | AN | Bergey's Manual | Based on Prevotellaceae |
| Prevotellaceae | Prevotella_7 | A | Bergey's Manual | Based on Prevotella |
| Prevotellaceae | Prevotella_9 | A | NA | |
| Prevotellaceae | Prevotellaceae UCG-001 | A | Bergey's Manual | Based on Prevotellaceae |
| Prevotellaceae | Prevotellaceae UCG-004 | A | Bergey's Manual | Based on Prevotellaceae |
| Promicromonosporaceae | Isoptericola | A | Bergey's Manual | |
| Propionibacteriaceae | Friedmanniella | A | Bergey's Manual | |
| Propionibacteriaceae | Marinilutecoccus | N/A | | |
| Propionibacteriaceae | Microlunatus | A | Bergey's Manual | |
| Propionibacteriaceae | Other | N/A | NA | |
| Propionibacteriaceae | Tessaracoccus | A | Bergey's Manual | |
| Pseudomonadaceae | Pseudomonas | A | Bergey's Manual | |
| Pseudonocardiaceae | Actinomycetospora | A | Bergey's Manual | |
| Pseudonocardiaceae | Actinophytocola | A | Bergey's Manual | Based on Pseudonocardiaceae |
| Pseudonocardiaceae | Crossiella | A | Bergey's Manual | Based on Pseudonocardiaceae |
| Pseudonocardiaceae | Kibdelosporangium | A | Bergey's Manual | |
| Pseudonocardiaceae | Other | A | Bergey's Manual | Based on Pseudonocardiaceae |
| Pseudonocardiaceae | Pseudonocardia | A | Bergey's Manual | |

| | | | | |
|---|---|---|---|---|
| Puniceicoccaceae | Cerasicoccus | A | Bergey's Manual | |
| Reyranellaceae | Reyranella | N/A | | |
| Rhizobiaceae | Ahrensia | A | Bergey's Manual | |
| Rhizobiaceae | Aliihoeflea | A | Bergey's Manual | Based on Rhizobiaceae |
| Rhizobiaceae | Allorhizobium-Neorhizob | A | Bergey's Manual | Based on Rhizobium and Allorhizobi |
| Rhizobiaceae | Aminobacter | A | Bergey's Manual | |
| Rhizobiaceae | Aquamicrobium | N/A | Bergey's Manual | |
| Rhizobiaceae | Aurantimonas | A | Bergey's Manual | Based on Rhizobiaceae |
| Rhizobiaceae | Aureimonas | A | Bergey's Manual | Based on Rhizobiaceae |
| Rhizobiaceae | Brucella | A | Bergey's Manual | |
| Rhizobiaceae | Corticibacterium | A | Bergey's Manual | Based on Rhizobiaceae |
| Rhizobiaceae | Falsochrobactrum | N/A | Bergey's Manual | |
| Rhizobiaceae | Hoeflea | A | Bergey's Manual | Based on Rhizobiaceae |
| Rhizobiaceae | Jiella | A | Bergey's Manual | Based on Rhizobiaceae |
| Rhizobiaceae | Mesorhizobium | A | Bergey's Manual | |
| Rhizobiaceae | Neorhizobium | A | Bergey's Manual | Based on Rhizobiaceae |
| Rhizobiaceae | Ochrobactrum | A | Bergey's Manual | |
| Rhizobiaceae | Other | A | Bergey's Manual | Based on Rhizobiaceae |
| Rhizobiaceae | Other | A | Bergey's manual | |
| Rhizobiaceae | Paenochrobactrum | A | Bergey's Manual | |
| Rhizobiaceae | Phyllobacterium | A | Bergey's Manual | |
| Rhizobiaceae | Pseudaminobacter | A | Bergey's Manual | |
| Rhizobiaceae | Shinella | A | Bergey's Manual | Based on Rhizobiaceae |
| Rhizobiaceae | Tianweitania | A | Bergey's Manual | Based on Rhizobiaceae |
| Rhodanobacteraceae | Chujaibacter | N/A | | |
| Rhodanobacteraceae | Dokdonella | N/A | | |
| Rhodanobacteraceae | Luteibacter | N/A | | |
| Rhodanobacteraceae | Mizugakiibacter | N/A | | |
| Rhodanobacteraceae | Oleiagrimonas | N/A | | |
| Rhodanobacteraceae | Rhodanobacter | A | Bergey's Manual | |
| Rhodobacteraceae | Actibacterium | A | Bergey's Manual | Based on Rhodobacteraceae |
| Rhodobacteraceae | Albirhodobacter | A | Bergey's Manual | Based on Rhodobacteraceae |
| Rhodobacteraceae | Amaricoccus | A | Bergey's Manual | |
| Rhodobacteraceae | Boseongicola | A | Bergey's Manual | Based on Rhodobacteraceae |
| Rhodobacteraceae | Defluviimonas | A | Bergey's Manual | Based on Rhodobacteraceae |
| Rhodobacteraceae | Falsirhodobacter | A | Bergey's Manual | Based on Rhodobacteraceae |
| Rhodobacteraceae | Gemmobacter | A | Bergey's Manual | |
| Rhodobacteraceae | Jannaschia | A | Bergey's Manual | Based on Rhodobacteraceae |
| Rhodobacteraceae | Limibaculum | A | Bergey's Manual | Based on Rhodobacteraceae |
| Rhodobacteraceae | Maribius | A | Bergey's Manual | Based on Rhodobacteraceae |
| Rhodobacteraceae | Oceaniovalibus | A | Bergey's Manual | Based on Rhodobacteraceae |

| | | | | |
|---|---|---|---|---|
| Rhodobacteraceae | Octadecabacter | A | Bergey's Manual | |
| Rhodobacteraceae | Other | A | Bergey's Manual | Based on Rhodobacteraceae |
| Rhodobacteraceae | Other | A | Bergey's manual | |
| Rhodobacteraceae | Paenirhodobacter | A | Bergey's Manual | Based on Rhodobacteraceae |
| Rhodobacteraceae | Paracoccus | A | Bergey's Manual | |
| Rhodobacteraceae | Plastorhodobacter | A | Bergey's Manual | Based on Rhodobacteraceae |
| Rhodobacteraceae | Pseudorhodobacter | A | Bergey's Manual | Based on Rhodobacteraceae |
| Rhodobacteraceae | Pseudoruegeria | A | Bergey's Manual | Based on Rhodobacteraceae |
| Rhodobacteraceae | Rhodobacter | N/A | Bergey's Manual | |
| Rhodobacteraceae | Rhodobaculum | A | Bergey's Manual | Based on Rhodobacteraceae |
| Rhodobacteraceae | Roseivivax | A | Bergey's Manual | |
| Rhodobacteraceae | Roseovarius | A | Bergey's Manual | |
| Rhodobacteraceae | Rubellimicrobium | A | Bergey's Manual | Based on Rhodobacteraceae |
| Rhodobacteraceae | Sulfitobacter | A | Bergey's Manual | |
| Rhodobacteraceae | Thioclava | A | Bergey's Manual | Based on Rhodobacteraceae |
| Rhodobacteraceae | Yoonia-Loktanella | A | Bergey's Manual | Based on Yoonia |
| Rhodocyclaceae | Azovibrio | A | Bergey's Manual | |
| Rhodocyclaceae | Thauera | A | Bergey's Manual | |
| Rhodomicrobiaceae | Rhodomicrobium | A | Bergey's Manual | |
| Rhodothermaceae | Rubrivirga | A | Bergey's Manual | Based on Rhodothermaceae |
| Rickettsiaceae | Rickettsia | N/A | Bergey's Manual | |
| Rikenellaceae | Alistipes | AN | Bergey's Manual | |
| Rikenellaceae | Other | N/A | | |
| Rikenellaceae | Other | N/A | | |
| Rikenellaceae | Rikenella | AN | Bergey's Manual | |
| Rikenellaceae | Rikenellaceae RC9 gut gr | A | Bergey's Manual | Based on Rikenellaceae |
| Rs-E47 termite group | Other | N/A | NA | |
| Rs-E47 termite group | Other | N/A | | |
| Rubinisphaeraceae | Planctomicrobium | N/A | | |
| Rubinisphaeraceae | SH-PL14 | N/A | | |
| Rubritaleaceae | Luteolibacter | A | Bergey's Manual | Based on Rubritaleaceae |
| Ruminococcaceae | [Eubacterium] siraeum g | AN | Bergey's Manual | Based on Eubacterium |
| Ruminococcaceae | Anaerotruncus | AN | Bergey's Manual | |
| Ruminococcaceae | Angelakisella | AN | Bergey's Manual | Based on Ruminococcaceae |
| Ruminococcaceae | Candidatus Soleaferrea | AN | Bergey's Manual | Based on Ruminococcaceae |
| Ruminococcaceae | Caproiciproducens | AN | Bergey's Manual | Based on Ruminococcaceae |
| Ruminococcaceae | DTU089 | AN | Bergey's Manual | Based on Ruminococcaceae |
| Ruminococcaceae | Faecalibacterium | AN | Bergey's Manual | |
| Ruminococcaceae | Fournierella | AN | Bergey's Manual | Based on Ruminococcaceae |
| Ruminococcaceae | Harryflintia | AN | Bergey's Manual | Based on Ruminococcaceae |
| Ruminococcaceae | Incertae Sedis | AN | Bergey's Manual, Browne et al., 2( | Based on Ruminococcaceae |

| | | | | |
|---|---|---|---|---|
| Ruminococcaceae | Negativibacillus | AN | Bergey's Manual | Based on Ruminococcaceae |
| Ruminococcaceae | Other | AN | Bergey's Manual | Based on Ruminococcaceae |
| Ruminococcaceae | Other | AN | Bergey's manual | |
| Ruminococcaceae | Paludicola | AN | Bergey's Manual | Based on Ruminococcaceae |
| Ruminococcaceae | Pygmaiobacter | AN | Bergey's Manual | Based on Ruminococcaceae |
| Ruminococcaceae | Ruminococcus | AN | Bergey's Manual, Browne et al., 2016 | |
| Ruminococcaceae | Subdoligranulum | AN | Bergey's Manual | |
| Ruminococcaceae | UBA1819 | AN | Bergey's Manual | Based on Faecalibacterium |
| Salinisphaeraceae | Salinisphaera | A | Bergey's Manual | |
| Sandaracinaceae | Other | N/A | NA | |
| Sanguibacteraceae | Sanguibacter-Flavimobili | N/A | Bergey's Manual | Based on Sanguibacter |
| Saprospiraceae | Lewinella | A | Bergey's Manual | Based on Saprospiraceae |
| Saprospiraceae | Other | A | Bergey's Manual | Based on Saprospiraceae |
| Saprospiraceae | Other | A | Bergey's manual | |
| SC-I-84 | Other | N/A | NA | |
| SC-I-84 | Other | N/A | | |
| Schlesneriaceae | Planctopirus | N/A | | |
| Schlesneriaceae | Schlesneria | A | Bergey's Manual | |
| Shewanellaceae | Shewanella | A | Bergey's Manual | |
| Solirubrobacteraceae | Conexibacter | A | Bergey's Manual | |
| Solirubrobacteraceae | Other | N/A | Bergey's Manual | Based on Solirubrobacteraceae |
| Solirubrobacteraceae | Parviterribacter | N/A | NA | |
| Solirubrobacteraceae | Patulibacter | A | Bergey's Manual | |
| Solirubrobacteraceae | Solirubrobacter | A | Bergey's Manual | |
| Sphingobacteriaceae | Pedobacter | A | Bergey's Manual | |
| Sphingobacteriaceae | Sphingobacterium | A | Bergey's Manual | |
| Sphingomonadaceae | Altererythrobacter | N/A | | |
| Sphingomonadaceae | Erythrobacter | A | Bergey's Manual | |
| Sphingomonadaceae | Novosphingobium | A | Takeuchi et al., 2001, Bergey's manual | |
| Sphingomonadaceae | Qipengyuania | N/A | | |
| Sphingomonadaceae | Sphingomonas | A | Bergey's Manual | |
| Sphingomonadaceae | Sphingopyxis | A | Bergey's Manual | |
| Spirosomaceae | Persicitalea | N/A | | |
| Spirosomaceae | Rhabdobacter | N/A | | |
| Sporichthyaceae | Longivirga | A | Bergey's Manual | Based on Sporichthyaceae |
| Sporichthyaceae | Other | A | Bergey's Manual | Based on Sporichthyaceae |
| Sporomusaceae | Dendrosporobacter | AN | Bergey's Manual | |
| Sporomusaceae | Other | N/A | NA | |
| Sporomusaceae | Other | N/A | | |
| Sporomusaceae | Sporomusa | A | Bergey's Manual | |
| Staphylococcaceae | Corticicoccus | N/A | | |

| | | | | |
|---|---|---|---|---|
| Staphylococcaceae | Jeotgalicoccus | A | Bergey's Manual | |
| Staphylococcaceae | Macrococcus | N/A | | |
| Staphylococcaceae | Other | N/A | NA | |
| Staphylococcaceae | Other | N/A | | |
| Staphylococcaceae | Staphylococcus | A | Bergey's Manual | |
| Stappiaceae | Labrenzia | N/A | | |
| Stappiaceae | Other | N/A | NA | |
| Streptococcaceae | Lactococcus | A | Bergey's Manual | |
| Streptococcaceae | Other | A | Bergey's Manual | Based on Streptococcaceae |
| Streptococcaceae | Other | A | Bergey's manual | |
| Streptococcaceae | Streptococcus | A | Bergey's Manual | |
| Streptomycetaceae | Kitasatospora | A | Bergey's Manual | Based on Streptomycetaceae |
| Streptomycetaceae | Other | A | Bergey's Manual | Based on Streptomycetaceae |
| Streptomycetaceae | Streptomyces | A | Bergey's Manual | |
| Streptosporangiaceae | Streptosporangium | A | Bergey's Manual | |
| Sulfobacillaceae | Other | N/A | NA | |
| Sulfobacillaceae | Other | N/A | | |
| Sumerlaeaceae | Sumerlaea | N/A | | |
| Sutterellaceae | Other | N/A | NA | |
| Sutterellaceae | Other | N/A | | |
| Sutterellaceae | Parasutterella | A | Nagai et al., 2009 | |
| Sutterellaceae | Sutterella | A | Bergey's Manual | |
| Tannerellaceae | Candidatus Vestibaculum | N/A | | |
| Tannerellaceae | Macellibacteroides | N/A | | |
| Tannerellaceae | Other | N/A | NA | |
| Tannerellaceae | Other | N/A | | |
| Tannerellaceae | Parabacteroides | AN | Sakamoto & Benno, 2006 | |
| Thermoactinomycetaceae | Other | A | Bergey's Manual | Based on Thermoactinomycetaceae |
| Thermoactinomycetaceae | Other | A | Bergey's manual | |
| Thermoactinomycetaceae | Risungbinella | A | Bergey's Manual | Based on Thermoactinomycetaceae |
| Trueperaceae | Truepera | A | Bergey's Manual | |
| Tsukamurellaceae | Tsukamurella | A | Bergey's Manual | |
| UCG-010 | Other | N/A | NA | |
| UCG-010 | Other | N/A | | |
| Vagococcaceae | Vagococcus | A | Bergey's Manual | |
| Veillonellaceae | Dialister | AN | Bergey's Manual | |
| Veillonellaceae | Veillonella | A | Bergey's Manual | |
| Vibrionaceae | Vibrio | A | Bergey's Manual | |
| WD2101 soil group | Other | N/A | NA | |
| Weeksellaceae | Candidatus Hemobacteriu | N/A | | |
| Weeksellaceae | Chishuiella | N/A | | |

| | | | |
|---|---|---|---|
| Weeksellaceae | Chryseobacterium | A | Bergey's Manual |
| Weeksellaceae | Empedobacter | A | Bergey's Manual |
| Wohlfahrtiimonadaceae | Ignatzschineria | N/A | |
| Wohlfahrtiimonadaceae | Wohlfahrtiimonas | N/A | |
| Xanthobacteraceae | Afipia | A | NA |
| Xanthobacteraceae | Bradyrhizobium | A | Bergey's Manual |
| Xanthobacteraceae | Other | N/A | NA |
| Xanthobacteraceae | Other | N/A | |
| Xanthobacteraceae | Pseudolabrys | N/A | |
| Xanthobacteraceae | Pseudorhodoplanes | N/A | |
| Xanthobacteraceae | Rhodoplanes | A | Bergey's Manual |
| Xanthobacteraceae | Rhodopseudomonas | A | Bergey's Manual |
| Xanthomonadaceae | Luteimonas | A | Bergey's Manual |
| Xanthomonadaceae | Lysobacter | A | Bergey's Manual |
| Xanthomonadaceae | Pseudoxanthomonas | A | Bergey's Manual |
| Xanthomonadaceae | SN8 | N/A | |
| Xanthomonadaceae | Stenotrophomonas | A | Bergey's Manual |
| Xanthomonadaceae | Thermomonas | A | Bergey's Manual |
| Xenococcaceae | Pleurocapsa PCC-7319 | A | Bergey's Manual |
| Yersiniaceae | Other | N/A | NA |
| Yersiniaceae | Other | N/A | |
| Yersiniaceae | Rahnella | A | Bergey's Manual |
| Yersiniaceae | Serratia | A | Bergey's Manual |
| Yersiniaceae | Yersinia | A | Bergey's Manual |
