## Supplementary material for "Early-life gut microbiota assembly patterns are conserved between laboratory and wild mice": References for Supplementary Table 1


Benga, L., Sager, M., & Christensen, H. (2018). From the [Pasteurella] pneumotropica complex to Rodentibacter spp.: an update on [Pasteurella] pneumotropica. In *Veterinary Microbiology* (Vol. 217, pp. 121–134). Elsevier B.V. https://doi.org/10.1016/j.vetmic.2018.03.011

Browne, H. P., Forster, S. C., Anonye, B. O., Kumar, N., Neville, B. A., Stares, M. D., Goulding, D., & Lawley, T. D. (2016). Culturing of "unculturable" human microbiota reveals novel taxa and extensive sporulation. *Nature*, *533*(7604), 543–546. https://doi.org/10.1038/nature17645

Cai, S., & Dong, X. (2010). Cellulosilyticum ruminicola gen. nov., sp. nov., isolated from the rumen of yak, and reclassification of Clostridium lentocellum as Cellulosilyticum lentocellum comb. nov. *International Journal of Systematic and Evolutionary Microbiology*, *60*(4), 845–849. https://doi.org/10.1099/ijs.0.014712-0

Chang DH, Rhee MS, Ahn S, et al. Faecalibaculum rodentium gen. nov., sp. nov., isolated from the faeces of a laboratory mouse [published correction appears in Antonie Van Leeuwenhoek. 2016 Mar;109(3):481]. *Antonie Van Leeuwenhoek*. 2015;108(6):1309-1318. doi:10.1007/s10482-015-0583-3

Clavel, T., Charrier, C., Wenning, M., & Haller, D. (2013). Parvibacter caecicola gen. nov., sp. nov., a bacterium of the family Coriobacteriaceae isolated from the caecum of a mouse. *International Journal of Systematic and Evolutionary Microbiology*, *63*(PART7), 2642–2648. https://doi.org/10.1099/ijs.0.045344-0

Clavel, T., Duck, W., Charrier, C., Wenning, M., Elson, C., & Haller, D. (2010). Enterorhabdus caecimuris sp. nov., a member of the family Coriobacteriaceae isolated from a mouse model of spontaneous colitis, and emended description of the genus Enterorhabdus Clavel et al. 2009. In *International Journal of Systematic and Evolutionary Microbiology* (Vol. 60, Issue 7, pp. 1527– 1531). Microbiology Society. https://doi.org/10.1099/ijs.0.015016-0

Cox LM, Sohn J, Tyrrell KL, et al. Description of two novel members of the family Erysipelotrichaceae: Ileibacterium valens gen. nov., sp. nov. and Dubosiella newyorkensis, gen. nov., sp. nov., from the murine intestine, and emendation to the description of Faecalibaculum rodentium [published correction appears in Int J Syst Evol Microbiol. 2017 Oct;67(10 ):4289]. *Int J Syst Evol Microbiol*. 2017;67(5):1247-1254. doi:10.1099/ijsem.0.001793

Degelmann DM, Kolb S, Dumont M, Murrell JC, Drake HL. Enterobacteriaceae facilitate the anaerobic degradation of glucose by a forest soil. *FEMS Microbiol Ecol*. 2009;68(3):312-319. doi:10.1111/j.1574-6941.2009.00681.x

Garcia R, Stadler M, Gemperlein K, Müller R. *Aetherobacter fasciculatus* gen. nov., sp. nov. and *Aetherobacter rufus* sp. nov., novel myxobacteria with promising biotechnological applications. *Int J Syst Evol Microbiol*. 2016;66(2):928-938. doi:10.1099/ijsem.0.000813

Gerritsen, J., Hornung, B., Ritari, J., Paulin, L., Rijkers, G., Schaap, P., de Vos, W., & Smidt, H. (2019). A comparative and functional genomics analysis of the genus Romboutsia provides insight into adaptation to an intestinal lifestyle. *BioRxiv*, 845511. https://doi.org/10.1101/845511

Hardham JM, King KW, Dreier K, et al. Transfer of Bacteroides splanchnicus to Odoribacter gen. nov. as Odoribacter splanchnicus comb. nov., and description of Odoribacter denticanis sp. nov., isolated from the crevicular spaces of canine periodontitis patients. *Int J Syst Evol Microbiol*. 2008;58(Pt 1):103-109. doi:10.1099/ijs.0.63458-0

Iino, T., Mori, K., Tanaka, K., Suzuki, K. I., & Harayama, S. (2007). Oscillibacter valericigenes gen. nov., sp. nov., a valerate-producing anaerobic bacterium isolated from the alimentary canal of a Japanese corbicula clam. *International Journal of Systematic and Evolutionary Microbiology*, *57*(8), 1840–1845. https://doi.org/10.1099/ijs.0.64717-0

Jeong, H., Lim, Y. W., Yi, H., Sekiguchi, Y., Kamagata, Y., & Chun, J. (2007). Anaerosporobacter mobilis gen. nov., sp. nov., isolated from forest soil. *International Journal of Systematic and Evolutionary Microbiology*, *57*(8), 1784–1787. https://doi.org/10.1099/ijs.0.63283-0

Johnson DB, Stallwood B, Kimura S, Hallberg KB. Isolation and characterization of Acidicaldus organivorus, gen. nov., sp. nov.: a novel sulfur-oxidizing, ferric iron-reducing thermo-acidophilic heterotrophic Proteobacterium. *Arch Microbiol*. 2006;185(3):212-221. doi:10.1007/s00203-006-0087-7

Kanso, S., & Patel, B. K. C. (2003). Microvirga subterranea gen. nov., sp. nov., a moderate thermophile from a deep subsurface Australian thermal aquifer. *International Journal of Systematic and Evolutionary Microbiology*, *53*(2), 401–406. https://doi.org/10.1099/ijs.0.02348- 0

Kim, B. C., Jeon, B. S., Kim, S., Kim, H., Um, Y., & Sang, B. I. (2015). Caproiciproducens galactitolivorans gen. Nov., sp. nov., a bacterium capable of producing caproic acid from galactitol, isolated from a wastewater treatment plant. *International Journal of Systematic and Evolutionary Microbiology*, *65*(12), 4902–4908. https://doi.org/10.1099/ijsem.0.000665

Kläring, K., Hanske, L., Bui, N., Charrier, C., Blaut, M., Haller, D., Plugge, C. M., & Clavel, T. (2013). Intestinimonas butyriciproducens gen. nov., sp. nov., a butyrate-producing bacterium from the mouse intestine. *International Journal of Systematic and Evolutionary Microbiology*, *63*(PART 12), 4606–4612. https://doi.org/10.1099/ijs.0.051441-0

Krishnamurthi, S., Bhattacharya, A., Mayilraj, S., Saha, P., Schumann, P., & Chakrabarti, T. (2009). Description of Paenisporosarcina quisquiliarum gen. nov., sp. nov., and reclassification of Sporosarcina macmurdoensis Reddy et al. 2003 as Paenisporosarcina macmurdoensis comb. nov. *International Journal of Systematic and Evolutionary Microbiology*, *59*(6), 1364–1370. https://doi.org/10.1099/ijs.0.65130-0

Lagkouvardos, I., Lesker, T. R., Hitch, T. C. A., Gálvez, E. J. C., Smit, N., Neuhaus, K., Wang, J., Baines, J. F., Abt, B., Stecher, B., Overmann, J., Strowig, T., & Clavel, T. (2019). Sequence and cultivation study of Muribaculaceae reveals novel species, host preference, and functional potential of this yet undescribed family. *Microbiome*, *7*(1), 28. https://doi.org/10.1186/s40168- 019-0637-2

Lee H, Jung KB, Kwon O, et al. *Limosilactobacillus reuteri* DS0384 promotes intestinal epithelial maturation via the postbiotic effect in human intestinal organoids and infant mice. *Gut Microbes*. 2022;14(1):2121580. doi:10.1080/19490976.2022.2121580

Liu X, Mao B, Gu J, et al. *Blautia*-a new functional genus with potential probiotic properties?. *Gut Microbes*. 2021;13(1):1-21. doi:10.1080/19490976.2021.1875796

Mediannikov, O., Sekeyová, Z., Birg, M.-L., & Raoult, D. (2010). A Novel Obligate Intracellular Gamma-Proteobacterium Associated with Ixodid Ticks, Diplorickettsia massiliensis, Gen. Nov., Sp. Nov. *PLoS ONE*, *5*(7), e11478. https://doi.org/10.1371/journal.pone.0011478

Morotomi M, Nagai F, Watanabe Y. Description of Christensenella minuta gen. nov., sp. nov., isolated from human faeces, which forms a distinct branch in the order Clostridiales, and proposal of Christensenellaceae fam. nov. *Int J Syst Evol Microbiol*. 2012;62(Pt 1):144-149. doi:10.1099/ijs.0.026989-0

Mozota M, Castro I, Gómez-Torres N, et al. Administration of *Ligilactobacillus salivarius* MP101 in an Elderly Nursing Home during the COVID-19 Pandemic: Immunological and Nutritional Impact. *Foods*. 2021;10(9):2149. Published 2021 Sep 11. doi:10.3390/foods10092149

Nagai, F., Morotomi, M., Sakon, H., & Tanaka, R. (2009). Parasutterella excrementihominis gen. nov., sp. nov., a member of the family Alcaligenaceae isolated from human faeces. *International Journal of Systematic and Evolutionary Microbiology*, *59*(7), 1793–1797. https://doi.org/10.1099/ijs.0.002519-0

Pfeiffer, N., Desmarchelier, C., Blaut, M., Daniel, H., Haller, D., & Clavel, T. (2012). Acetatifactor muris gen. nov., sp. nov., a novel bacterium isolated from the intestine of an obese mouse. *Archives of Microbiology*, *194*(11), 901–907. https://doi.org/10.1007/s00203-012-0822-1

Ricaboni D, Mailhe M, Cadoret F, Vitton V, Fournier PE, Raoult D. *'Colidextribacter massiliensis'* gen. nov., sp. nov., isolated from human right colon. *New Microbes New Infect*. 2016;17:27-29. Published 2016 Nov 28. doi:10.1016/j.nmni.2016.11.023

Rosero JA, Killer J, Sechovcová H, et al. Reclassification of *Eubacterium rectale* (Hauduroy *et al.* 1937) Prévot 1938 in a new genus *Agathobacter* gen. nov. as *Agathobacter rectalis* comb. nov., and description of *Agathobacter ruminis* sp. nov., isolated from the rumen contents of sheep and cows. *Int J Syst Evol Microbiol*. 2016;66(2):768-773. doi:10.1099/ijsem.0.000788

Sakamoto M, Benno Y. Reclassification of Bacteroides distasonis, Bacteroides goldsteinii and Bacteroides merdae as Parabacteroides distasonis gen. nov., comb. nov., Parabacteroides goldsteinii comb. nov. and Parabacteroides merdae comb. nov. *Int J Syst Evol Microbiol*. 2006;56(Pt 7):1599-1605. doi:10.1099/ijs.0.64192-0

Sarma-Rupavtarm RB, Ge Z, Schauer DB, Fox JG, Polz MF. Spatial distribution and stability of the eight microbial species of the altered schaedler flora in the mouse gastrointestinal tract. *Appl Environ Microbiol*. 2004;70(5):2791-2800. doi:10.1128/AEM.70.5.2791-2800.2004

Schnupf, P., Gaboriau-Routhiau, V., Gros, M., Friedman, R., Moya-Nilges, M., Nigro, G., Cerf- Bensussan, N., & Sansonetti, P. J. (2015). Growth and host interaction of mouse segmented filamentous bacteria in vitro. *Nature*, *520*(7545), 99–103. https://doi.org/10.1038/nature14027

Takeuchi, M., Hamana, K., & Hiraishi, A. (2001). Proposal of the genus Sphingomonas sensu stricto and three new genera, Sphingobium, Novosphingobium and Sphingopyxis, on the basis of phylogenetic and chemotaxonomic analyses. *International Journal of Systematic and Evolutionary Microbiology*, *51*(4), 1405–1417. https://doi.org/10.1099/00207713-51-4-1405

Tindall BJ. The names *Hungateiclostridium*Zhang *et al.* 2018, *Hungateiclostridium thermocellum* (Viljoen *et al.* 1926) Zhang *et al.*2018, *Hungateiclostridium cellulolyticum*(Patel *et al.* 1980) Zhang *et al.* 2018, *Hungateiclostridium aldrichii* (Yang *et al.* 1990) Zhang *et al.* 2018, *Hungateiclostridium alkalicellulosi* (Zhilina *et al.* 2006) Zhang *et al.*2018, *Hungateiclostridium clariflavum*(Shiratori *et al.* 2009) Zhang *et al.* 2018, *Hungateiclostridium straminisolvens* (Kato *et al.* 2004) Zhang *et al.* 2018 and *Hungateiclostridium saccincola* (Koeck *et al.*2016) Zhang *et al.* 2018 contravene Rule 51b of the International Code of Nomenclature of Prokaryotes and require replacement names in the genus *Acetivibrio* Patel *et al.* 1980. *Int J Syst Evol Microbiol*. 2019;69(12):3927-3932. doi:10.1099/ijsem.0.003685

Trachsel, J., Humphrey, S., & Allen, H. K. (2018). Butyricicoccus porcorum sp. nov., a butyrate- producing bacterium from swine intestinal tract. *International Journal of Systematic and Evolutionary Microbiology*, *68*(5), 1737–1742. https://doi.org/10.1099/ijsem.0.002738

Vandamme PA, Peeters C, Cnockaert M, et al. Bordetella bronchialis sp. nov., Bordetella flabilis sp. nov. and Bordetella sputigena sp. nov., isolated from human respiratory specimens, and reclassification of Achromobacter sediminum Zhang et al. 2014 as Verticia sediminum gen. nov., comb. nov. *Int J Syst Evol Microbiol*. 2015;65(10):3674-3682. doi:10.1099/ijsem.0.000473

Würdemann D, Tindall BJ, Pukall R, et al. Gordonibacter pamelaeae gen. nov., sp. nov., a new member of the Coriobacteriaceae isolated from a patient with Crohn's disease, and reclassification of Eggerthella hongkongensis Lau et al. 2006 as Paraeggerthella hongkongensis gen. nov., comb. nov. *Int J Syst Evol Microbiol*. 2009;59(Pt 6):1405-1415. doi:10.1099/ijs.0.005900-0

Yutin, N., & Galperin, M. Y. (2013). A genomic update on clostridial phylogeny: Gram-negative spore formers and other misplaced clostridia. *Environmental Microbiology*, *15*(10), 2631–2641. https://doi.org/10.1111/1462-2920.12173

Zheng J, Wittouck S, Salvetti E, et al. A taxonomic note on the genus *Lactobacillus*: Description of 23 novel genera, emended description of the genus *Lactobacillus*Beijerinck 1901, and union of *Lactobacillaceae*and *Leuconostocaceae*. *Int J Syst Evol Microbiol*. 2020;70(4):2782-2858. doi:10.1099/ijsem.0.004107
